# Regulation of inflammation and protection against invasive pneumococcal infection by the long pentraxin PTX3

**DOI:** 10.1101/2022.04.14.488329

**Authors:** Rémi Porte, Rita Silva-Gomes, Charlotte Theroude, Raffaella Parente, Fatemeh Asgari, Marina Sironi, Fabio Pasqualini, Sonia Valentino, Rosanna Asselta, Camilla Recordati, Andrea Doni, Antonio Inforzato, Carlos Rodriguez-Gallego, Ignacio Obando, Elena Colino, Barbara Bottazzi, Alberto Mantovani

## Abstract

*Streptococcus pneumoniae* is a major pathogen in children, elderly subjects and immunodeficient patients. PTX3 is a fluid phase pattern recognition molecule (PRM) involved in resistance to selected microbial agents and in regulation of inflammation. The present study was designed to assess the role of PTX3 in invasive pneumococcal infection. In a murine model of invasive pneumococcal infection, PTX3 was strongly induced in non-hematopoietic (particularly, endothelial) cells. The IL-1β/MyD88 axis played a major role in regulation of the *Ptx3* gene expression. *Ptx3*^-/-^ mice were more susceptible to invasive pneumococcal infection. Although high concentrations of PTX3 had opsonic activity *in vitro*, no evidence of PTX3-enhanced phagocytosis was obtained *in vivo*. In contrast, *Ptx3*-deficient mice showed enhanced recruitment of neutrophils and inflammation. Using P-selectin deficient mice, we found that protection against pneumococcus was dependent upon PTX3-mediated regulation of neutrophil inflammation. In humans, PTX3 genetic polymorphisms were associated with invasive pneumococcal infections. Thus, this fluid phase PRM plays an important role in tuning inflammation and resistance against invasive pneumococcal infection.

## Introduction

*Streptococcus pneumoniae* (or pneumococcus) is a major leading cause of bacterial pneumonia, meningitis and sepsis in children, elders and immunodeficient patients. This pathogen is estimated to be responsible for most of the lower respiratory infections, causing around 1.18 million deaths annually worldwide (Troeger et al., 2018). Despite the widespread use of pneumococcal conjugate vaccines and antibiotic treatments, the combination of high carriage rate, ability to become pathogenic to the host, and genetic adaptability, make pneumococcus a significant cause of community- and hospital-acquired infections (Weiser et al., 2018). Since 2017, *S. pneumoniae* is classified as one of the 12 priority pathogens by World Health Organization. *S. pneumoniae* is a Gram-positive extracellular opportunistic pathogen which colonizes the respiratory mucosa of the upper respiratory tract. Depending on the virulence factors expressed by the pathogen and host factors, the disease can evolve to pneumococcal invasive infection, where pneumococcus invades the lower respiratory tract and translocates through the blood stream into the systemic compartment (Weiser et al., 2018). The introduction over the years of pneumococcal vaccines able to protect against a variable and increasing number of different serotypes has been able to reduce the impact of the infection in susceptible populations (Tin Tin Htar et al., 2019; Troeger et al., 2018). However, for some serotypes available vaccines confer limited protection only. In particular, it has been reported that the 13-valent pneumococcal conjugate vaccine (Tin Tin Htar et al., 2019) fails to reduce the risk of infection by serotype 3, which associated with a complicated disease course and increased risk of death (Weinberger et al., 2010).

As a first line of defense against respiratory pathogens, innate immune Pattern Recognition Molecules (PRMs) recognize microbial components and modulate immune response to control infections. Among conserved fluid phase PRMs, Pentraxin 3 (PTX3), is a member of the pentraxin family characterized by multifunctional properties, including regulation of innate immunity during infections (Garlanda et al., 2018). PTX3 is expressed by various hematopoietic and non-hematopoietic cells in response to microbial moieties and inflammatory cytokines (i.e IL-1β and TNF), and has been associated with the control of various infections by promoting different anti-microbial mechanisms. Indeed, PTX3 participates directly to the elimination of selected microorganisms by promoting phagocytosis, activating the complement cascade and as a component of Neutrophil Extracellular Traps (NET) (Daigo et al., 2012; Jaillon et al., 2014, 2007; Moalli et al., 2010; Porte et al., 2019). Furthermore PTX3 modulates tissue remodeling (Doni et al., 2015) and inflammation by tuning complement activation and P-selectin-dependent transmigration (Deban et al., 2010; Lech et al., 2013), both involved in neutrophil recruitment and in the evolution of respiratory tract infections (Quinton and Mizgerd, 2015).

In humans, PTX3 plasma levels increase in the context of inflammation and selected infectious diseases, including pneumococcal pathologies (i.e community-acquired pneumonia, ventilator associated pneumonia, pneumococcal exacerbated chronic obstructive pulmonary disease), correlating with the severity of the disease and predicting the risk of mortality (Bilgin et al., 2018; Kao et al., 2013; Mauri et al., 2014; Porte et al., 2019; Saleh et al., 2019; Shi et al., 2020; Siljan et al., 2019; Thulborn et al., 2017). Single nucleotide polymorphisms (SNPs) in the *PTX3* gene have been associated with patient susceptibility to respiratory infections (Brunel et al., 2018; Chiarini et al., 2010; Cunha et al., 2015, 2014; He et al., 2018; Olesen et al., 2007; Wójtowicz et al., 2015).

The involvement of PTX3 in the control of selected respiratory pathogens and in the modulation of infection prompted us to investigate the role of this molecule in the control of pneumococcal infections. In a murine model of invasive pneumococcal infection, we observed that PTX3 genetic deficiency is associated with higher susceptibility to infection and higher respiratory tract inflammation. We also observed that PTX3, mainly produced by stromal non-hematopoietic cells during pneumococcal infection, modulates neutrophil recruitment by dampening P-selectin dependent neutrophil migration. Hence, PTX3 plays a non-redundant role in the control of *S. pneumoniae* infection, modulating neutrophil associated respiratory tissue damage and pneumococcal systemic dissemination.

## Results

### PTX3 expression during Pneumococcal invasive infection

In order to define the relevance of PTX3 in pneumococcal respiratory disease, we first investigated whether the protein is induced during infection. Thus, we used a murine model of pneumococcal invasive infection induced by *S. pneumoniae* serotype 3. Mice were challenged intranasally with 5×10^4^ CFU and sacrificed at different time points. As already described, *S. pneumoniae* serotype 3 causes bacterial colonization of the respiratory tract, then disseminates through the blood circulation and infects other organs like the spleen, resulting in death within 3 to 4 days (Figure S1A-B) (de Porto et al., 2019). As early as 6h post-infection, we detected a local expression of PTX3 in the alveolar compartment near the pulmonary veins (Figure 1A-B). At 12h post-infection, we were able to detect the PTX3 specific staining by the endothelial cells in the area where we can appreciate inflammatory cells infiltration. This association was confirmed 24h post-infection, when a strong PTX3 staining was present near the recruitment site of inflammatory cells forming inflammatory foci (Figure 1A). The kinetic of PTX3 production was confirmed by the quantification of PTX3^+^ area (Figure 1B) and by analysis of mRNA in the lung (Figure S1C). Interestingly, local and systemic production of PTX3 was strongly induced by the infection during the disseminating phase (Figure 1C). During this invasive infection we observed that *Ptx3* was upregulated mainly in the lung, aorta and heart, while other organs like brain, kidneys and liver did not show higher *Ptx3* expression compared to the uninfected mice (Figure S1D).

**Figure 1.**
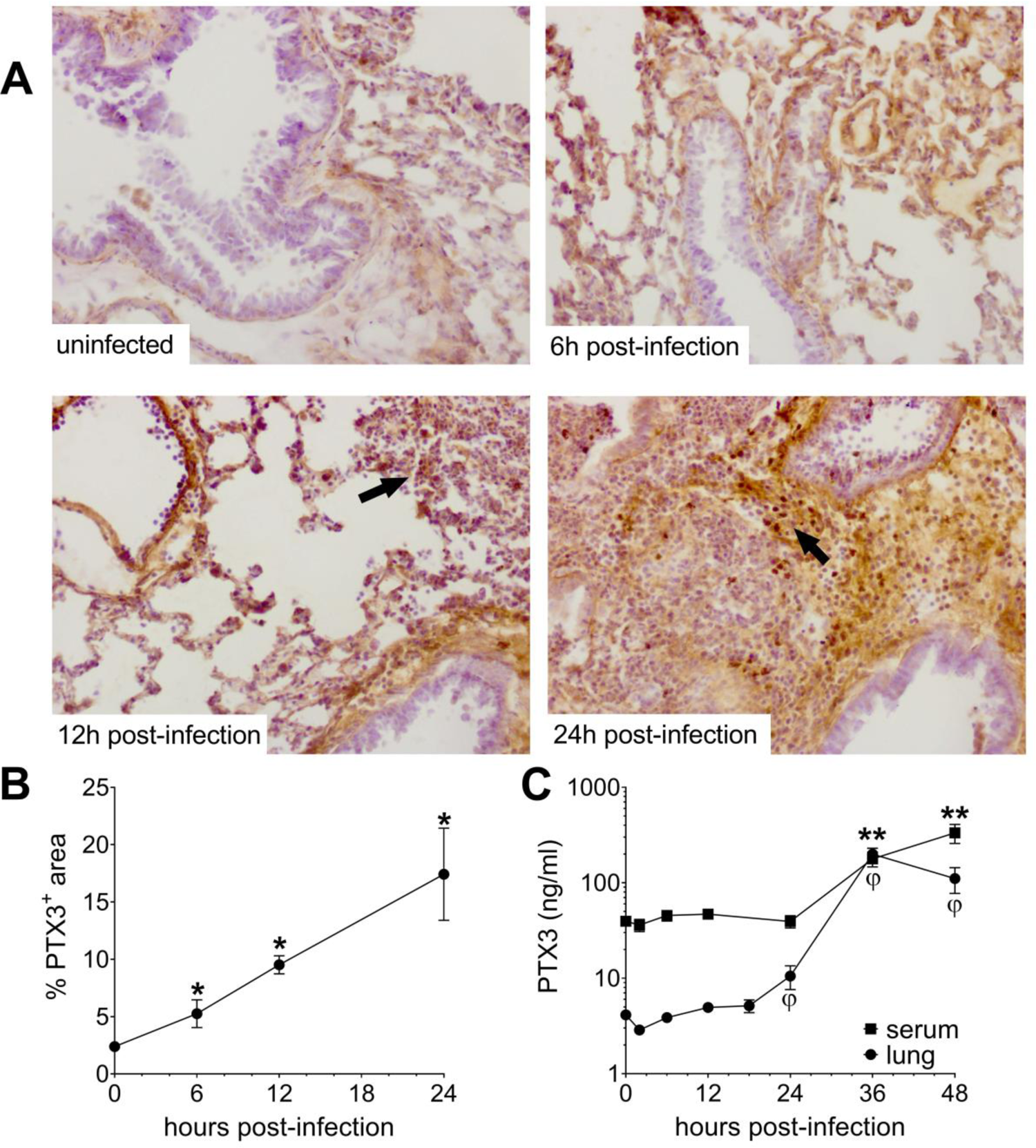
Invasive pneumococcal infection induces PTX3 expression. WT mice were infected intranasally with 5×10^4^ CFU of *S. pneumoniae* serotype 3 and sacrificed at the indicated time points for tissue collection. (A-B) Immunohistochemical analysis and quantification of PTX3 expression in lung sections (magnification ×10) from uninfected mice and mice sacrificed 6h, 12h and 24h post-infection (n=3-6). (A) One representative images of at least three biological replicates for each condition is reported. Inflammatory cell infiltrates are indicated by arrows. (B) Sections were scanned and analyzed to determine the percentage of PTX3^+^ area at the indicated time points. (C) PTX3 protein levels determined by ELISA in serum and lung homogenates collected at the indicated time points (n=4-10). Results are reported as mean ± SEM. Statistical significance was determined using the Mann-Whitney test comparing results to uninfected mice (φ or \**P*<0.05 and \*\**P*<0.01).

### Induction of PTX3 by IL-1β during *S. pneumoniae* infection

PTX3 has been described to be induced by primary inflammatory cytokines in particular IL-1β (Garlanda et al., 2018; Porte et al., 2019). In this pneumococcal invasive infection model we observed a rapid induction of IL-1 β (Figure 2A), and a strong correlation between the levels of IL-1β expressed in the respiratory tract with the levels of lung PTX3 (Figure 2B). Moreover, *Il1r*^-/-^ mice infected by *S. pneumoniae* showed lower PTX3 levels, locally and systemically (i.e in the lung and the serum respectively) (Figure 2C-D). *S. pneumoniae* infected *Myd88*^-/-^ mice were not able to produce PTX3 in the lung and presented the same impairment of PTX3 production as *Il1r*^-/-^ mice (Figure 2C-D). These data suggest that IL-1β is a major driver of PTX3 during pneumococcal infection.

**Figure 2.**
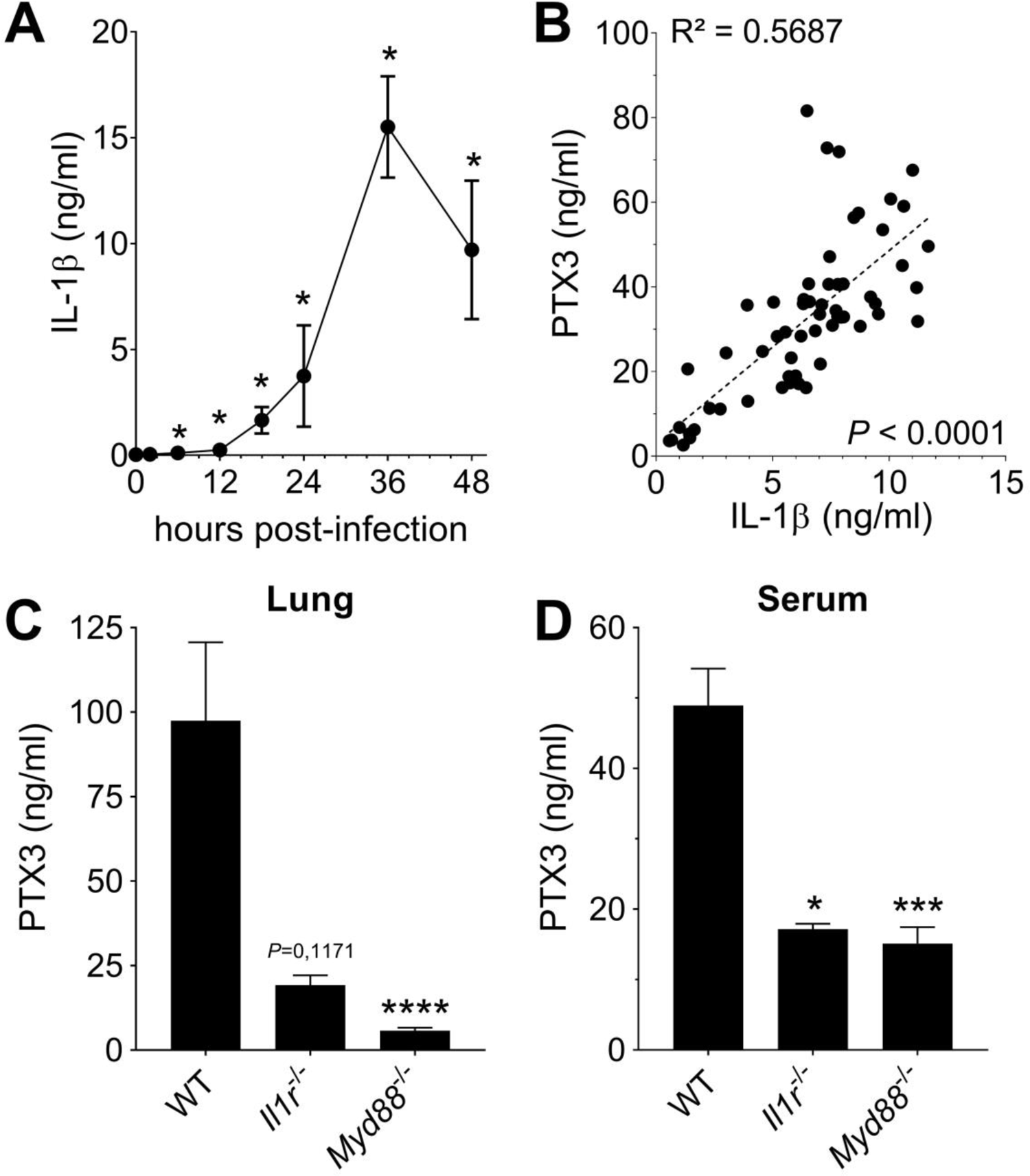
Role of IL-1β in induction of PTX3 during *S. pneumoniae* infection. WT mice were infected intranasally with 5×10^4^ CFU of *S. pneumoniae* serotype 3 and sacrificed at the indicated time points for tissue collection. (A) IL-1β protein levels in lung homogenates collected at the indicated time points determined by ELISA (n=3-4). (B) Correlation between PTX3 and IL-1β protein levels in lung homogenates of all infected mice sacrificed from 2h to 48h post-infection (data pooled from 5 independent experiments, n=60). PTX3 protein levels determined by ELISA in lung homogenates (C) and serum (D) collected 36h post-infection in WT, *Il1r*^-/-^ and *Myd88*^-/-^ mice (n=7-8). Results are reported as mean ± SEM. Statistical significance was determined using the Mann-Whitney test comparing results to uninfected mice (A-B) or the non-parametric Krukal-Wallis test with post-hoc corrected Dunn’s test comparing means to WT infected mice (C-D) (\**P*<0.05, \*\*\**P*<0.001 and \*\*\*\**P*<0.0001).

### Non-hematopoietic cells are a major source of PTX3 during pneumococcal infection

It has been previously reported that neutrophils contain preformed PTX3, representing an important source of the protein, rapidly released in response to proinflammatory cytokines or microbial recognition (Jaillon et al., 2007). In agreement, we observed that human neutrophils can release PTX3 upon stimulation with *S. pneumoniae* (Figure S2A). To investigate the involvement of neutrophils in the production of PTX3 in our model, we used mice lacking granulocyte colony-stimulating factor receptor (*Csf3r*^-/-^). These mice are characterized by chronic neutropenia, granulocyte and macrophage progenitor cell deficiency and impaired neutrophil mobilization (Liu et al., 1996; Ponzetta et al., 2019). Following pneumococcal infection, *Csf3r*^-/-^ mice presented lower levels of myeloperoxidase (MPO), a marker of neutrophil recruitment, in lung homogenates at 36h post-infection (Figure S2B). By contrast, even though these mice presented a lower amount of neutrophils recruited in response to the infection, they expressed the same pulmonary levels of PTX3 as WT mice (Figure S2B). These results suggest that neutrophils are not the main source of PTX3 in our murine model of pneumococcal invasive infection.

Since PTX3 can be produced by hematopoietic and non-hematopoietic cells, bone marrow chimeras were used to evaluate the cellular compartment responsible for PTX3 production. During pneumococcal infection, we did not observe any difference in the levels of PTX3 in the respiratory tract and in the serum of WT mice receiving bone marrow from *Ptx3^-/-^* or WT animals, while no PTX3 was measured in *Ptx3^-/-^* mice receiving WT or *Ptx3^-/-^* bone marrow (Figure 3A-B). These results suggest that PTX3 is mainly produced by the non-hematopoietic compartment after pneumococcal infection. Endothelial cells were described as an important source of PTX3 (Garlanda et al., 2018), thus we evaluated their contribution to PTX3 production during pneumococcal infection. To this aim we crossed conditional *Ptx3* deficient mice (*Ptx3LoxP^+/+^*) with *Cdh5-Cre* mice to generate animals with the deletion of PTX3 in endothelial cells. When *Ptx3LoxP^+/+^/Cdh5Cre^+/+^* mice were infected with *S. pneumoniae*, they presented approximately 50% reduction of PTX3 levels compared to PTX3-competent mice (Figure 3C-D). *In-vitro* experiments confirmed the ability of both murine and human endothelial cells to produce PTX3 after stimulation with *S. pneumoniae* (Figure S2C). Thus, in our setting, non-hematopoietic cells, mainly endothelial cells, are a major source of PTX3.

**Figure 3.**
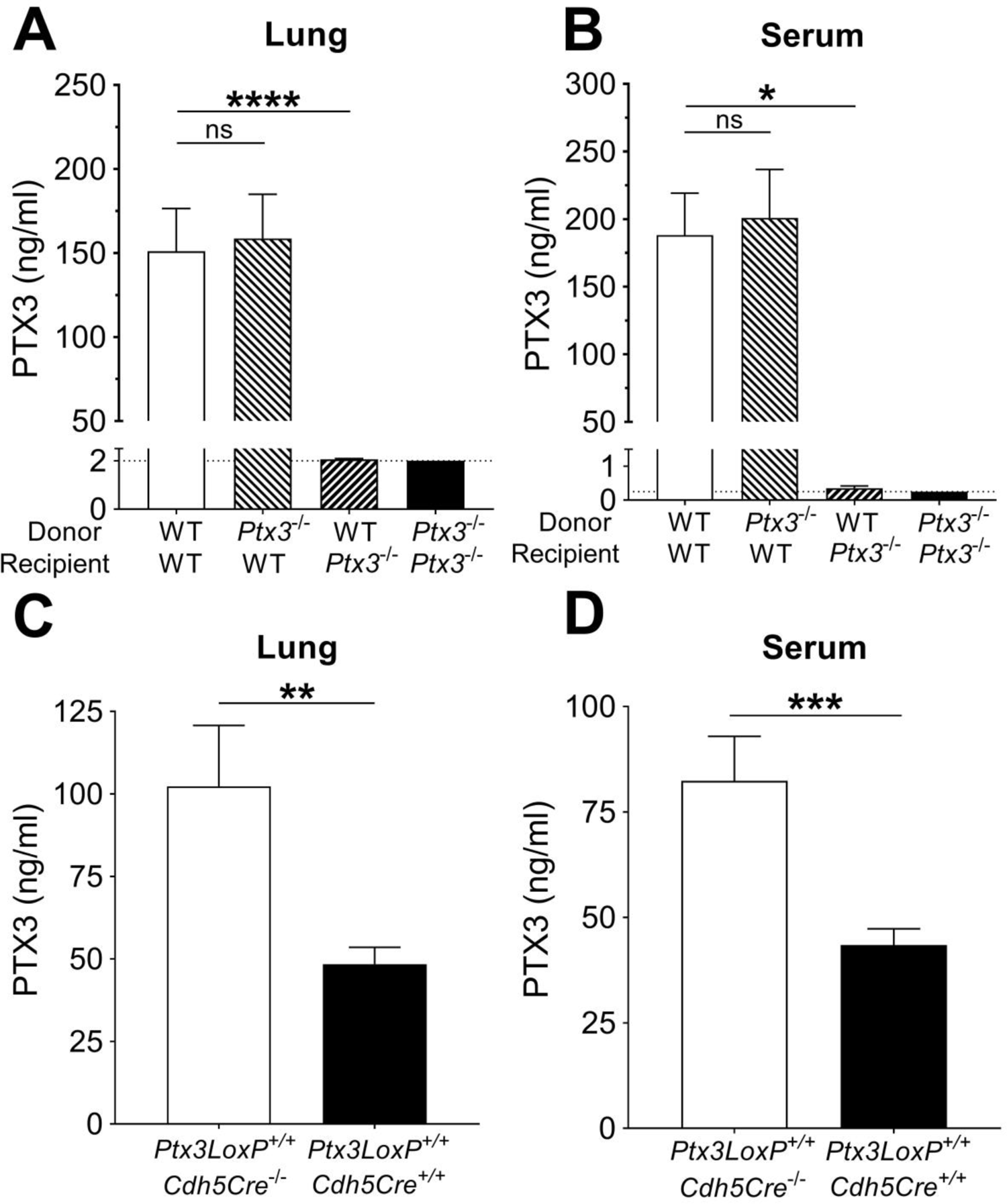
Non-hematopoietic cells are a major source of PTX3 during pneumococcal infection. Mice were infected intranasally with 5×10^4^ CFU of *S. pneumoniae* serotype 3 and sacrificed at 36h post-infection for tissue collection. (A-B) PTX3 protein levels determined by ELISA in lung homogenates (n=12-14, A) and serum (n=6, B) from chimeric mice. The experiment was repeated a second time with similar results. (C-D) PTX3 protein levels determined by ELISA in lung homogenates (C) and serum (D) collected from *Ptx3LoxP^+/+^Cdh5*-cre^-/-^, *Ptx3LoxP^+/+^Cdh5*-cre^+/+^ (n=10-13). Results are reported as mean ± SEM; PTX3 detection limit is 2 ng/ml in lung homogenates (A) and 0.25 ng/ml in serum (B) and is represented by a dotted line. Statistical significance was determined using the non-parametric Krukal-Wallis test with post-hoc corrected Dunn’s test comparing means to the WT recipient mice reconstituted with WT bone marrow (A-B) or the Mann-Whitney test (C-D) (\**P*<0.05, \*\**P*<0.01, \*\*\**P*<0.001 and **** *P*<0.0001).

### Non-redundant role of PTX3 in resistance to pneumococcal infection

Next we evaluated the role of PTX3 in resistance against pneumococcus. When *Ptx3*^-/-^ mice were infected with *S. pneumoniae* (5×10^4^ CFU), a significant increase of the bacterial load in the lung was observed during the invasive phase of infection (i.e. 36h post-infection), compared to WT mice (Figure 4A). Defective local control of bacterial growth was associated to an increase of bacterial load in the systemic compartment (Figure 4B). Interestingly there was no difference at earlier time points (i.e. 18h post-infection, Figure S3A), suggesting that PTX3 exerted a role in the control of pneumococcal infection mainly during the invasive phase. Using a bacterial dose (5×10^3^ CFU) inducing around 30% mortality in WT animals, *Ptx3*^-/-^ mice showed a significant higher mortality (83.3%; *P*<0.001) (Figure 4C). The phenotype described so far is not restricted to serotype 3 pneumococcus. In fact, when mice were infected with *S. pneumoniae* serotype 1, we observed a strong PTX3 production during the invasive phase of the infection (Figure S3B) and a correlation with IL-1β levels (Figure S3C). *Ptx3*^-/-^ mice infected by serotype 1 presented a higher sensitivity to the infection compared to WT animals, with a higher number of bacteria at the local site of infection and also in the systemic compartment 24h post-infection (Figure S3D-E). Thus, in the applied model of *S. pneumoniae* infection, the protection conferred by PTX3 is not limited to serotype 3, and embraces other bacterial serotypes of clinical relevance, including serotype 1.

**Figure 4.**
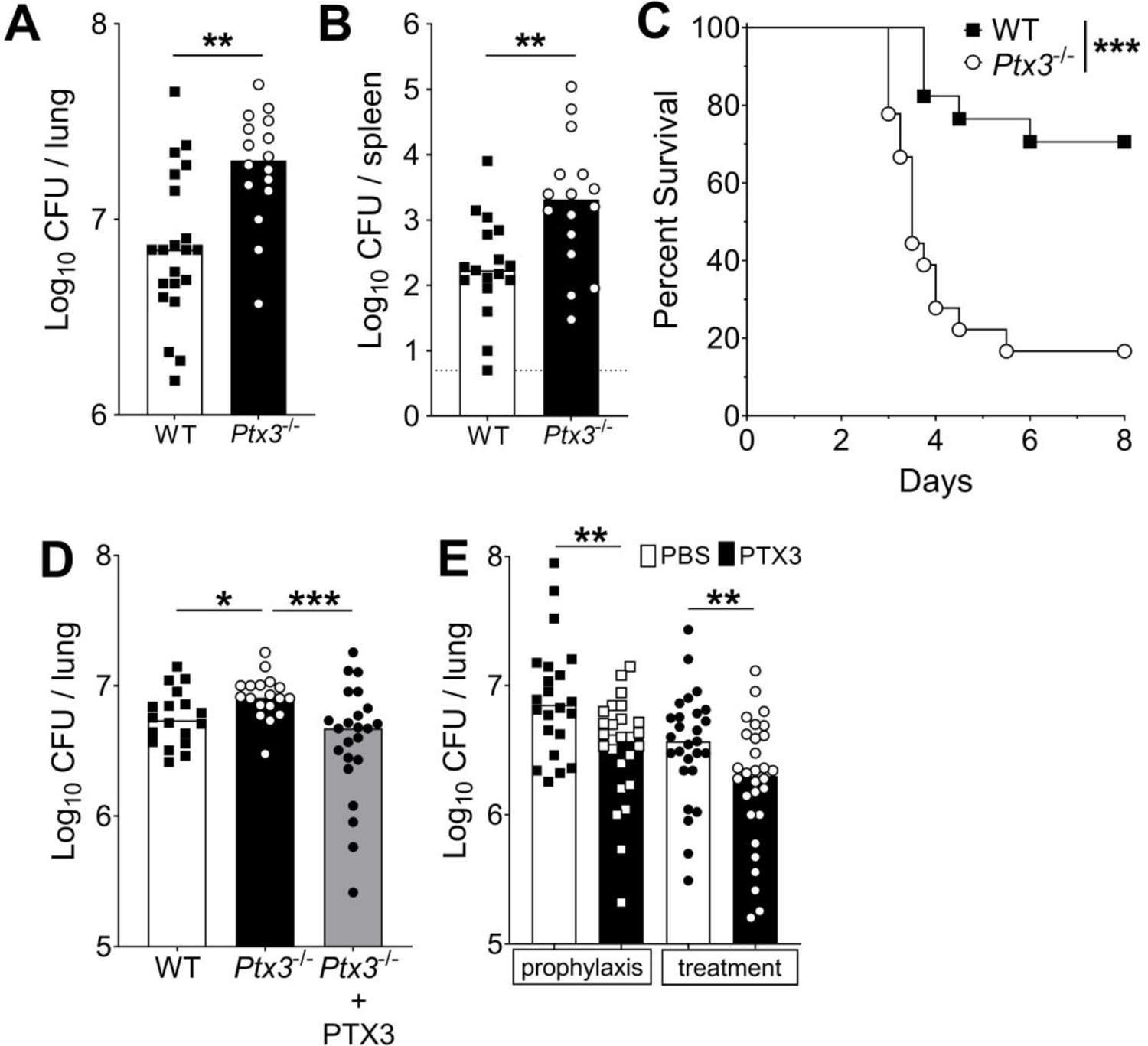
Defective resistance of PTX3-deficient mice to invasive pneumococcal infection. Mice were infected intranasally with different doses of *S. pneumoniae* serotype 3 and sacrificed at the indicated time points for tissue collection. (A-B) WT and *Ptx3*^-/-^ mice were infected with 5×10^4^ CFU and bacterial load in lung (A) and spleen (B) was analyzed at 36h post-infection (data pooled from 2 independent experiments, n=16-21). (C) Survival of WT and *Ptx3*^-/-^ mice (data pooled from 2 independent experiments, n=18) was monitored every 6h after infection with 5×10^3^ CFU. (D) Bacterial load was analyzed in lungs collected 36h post-infection from WT, *Ptx3*^-/-^ and *Ptx3*^-/-^ mice treated intraperitoneally with recombinant PTX3 (10µg/100µl) before the infection and 24h post-infection (n=18-23). (E) Bacterial load in lungs collected 36h post-infection from WT mice treated intranasally before the infection (prophylaxis, data pooled from 2 independent experiments, n=22-26) or 12h post-infection (treatment, data pooled from 3 independent experiments, n=37-40) with 1µg/30µl of recombinant PTX3 or PBS. Results are reported as median. CFU detection limits in the spleen is 5 CFU represented by a dotted line. Statistical significance was determined using the Mann-Whitney test (A-B, E), the non-parametric Krukal-Wallis test with post-hoc corrected Dunn’s test comparing means to the WT mice (D) and Log-rank (Mantel-Cox) test for survival (\**P*<0.05, \*\**P*<0.01 and \*\*\**P*<0.001).

Systemic administration of recombinant PTX3 to *Ptx3*^-/-^ mice rescues the phenotype. As reported in Figure 4D, PTX3 administration in *Ptx3*^-/-^ mice reduced lung colonization to the same level observed in WT mice. We then evaluated the antibacterial activity of PTX3 on *S. pneumoniae* serotype 3. WT animals were treated locally with 1µg of recombinant protein before infection or 12h post-infection. Under both conditions we observed a significant reduction (44% and 57% respectively; *P*<0.01) of the pulmonary bacterial load compared with the CFU found in mice treated with vehicle alone (Figure 4E).

### Lack of effective opsonic activity of PTX3

In an effort to explore the mechanism responsible for PTX3-mediated resistance, we first assessed the effect of the recombinant protein on the *in-vitro* growth of *S. pneumoniae*. The incubation of *S. pneumoniae* with 25-250 µg/ml of recombinant PTX3 did not have any effect on the growth rate of the bacteria (Figure S4).

PTX3 has the capability to act as an opsonin binding selected pathogens and increasing their removal by phagocytosis (Garlanda et al., 2002; Jaillon et al., 2014; Moalli et al., 2010). To assess whether the control of the pneumococcal infection by PTX3 was due to opsonic activity, we first analyzed PTX3 binding to *S. pneumoniae.* By using a flow cytometry assay, we analyzed PTX3 binding to *S. pneumoniae* serotype 3 mimicking the bacteria/PTX3 ratio found in the infected lung (10^6^ CFU/100 ng PTX3). Under these conditions, we did not observe any interaction of PTX3 with bacteria and, even with an amount of PTX3 5- to 10-fold higher than the one produced in the entire lung, less than 1% of the bacteria were bound (Figure 5A). At 500 µg/ml of PTX3 (5000-fold higher than in the lung homogenates) we observed binding to only 36.4% of bacteria (Figure 5A).

**Figure 5.**
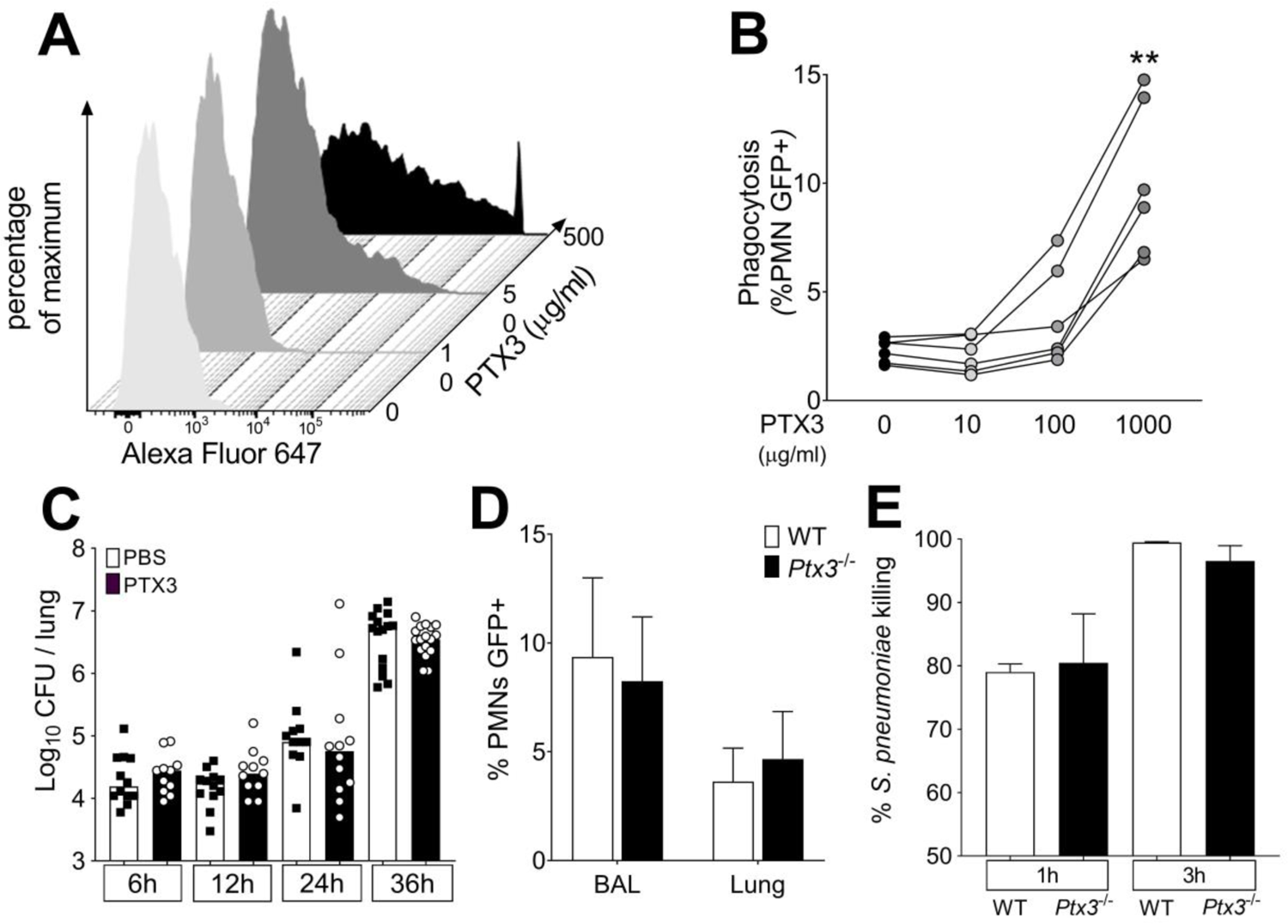
Role of phagocytosis in PTX3-mediated resistance to *S. pneumoniae*. (A) Binding of recombinant PTX3-biot at the indicated concentration with 10^6^ CFU of *S. pneumoniae* serotype 3. PTX3 binding to *S. pneumoniae* was analyzed by flow cytometry after incubation with Streptavidin-Alexa Fluor 647. (B) *S. pneumoniae* serotype 1 expressing GFP (10^6^ CFU) was pre-opsonized with indicated concentration of recombinant PTX3 and incubated 30 min with 10^5^ purified human neutrophils from 6 independent donors. GFP positive neutrophils were analyzed by flow cytometry. (C) Bacterial load in lung collected at indicated time points from WT mice infected intranasally with *S. pneumoniae* serotype 3 pre-opsonized with 33µg/ml of recombinant PTX3 or non-opsonized. (data pooled from 2 independent experiments, n=11-17). (D) Neutrophil phagocytosis of *S. pneumoniae* serotype 1 expressing GFP was analyze by flow cytometry. BAL and lungs from WT and *Ptx3*^-/-^ mice were collected 24h after infection with a lethal inoculum of *S. pneumoniae* (data pooled from 2 independent experiments, n=9-14). (E) alamarBlue based killing assay performed with Neutrophils purified from WT and *Ptx3^-/-^* mice assessed after 1h and 3h incubation at a MOI *S. pneumoniae*/Neutrophils 2/1. Results are expressed as mean of 5 technical replicate for each time point and donor (B), median (C) and mean ± SD (D-E). Statistical significance was determined using a one-way ANOVA with Sidak’s multiple comparison test (B), the non-parametric Krukal-Wallis test with post-hoc corrected Dunn’s test comparing means to the WT mice of each time point (C, E) and the Mann-Whitney test (D) (\*\**P*<0.01).

We then assessed phagocytosis *in vitro* and *in vivo* using GFP-expressing *S. pneumoniae* serotype 1 (*S. pneumoniae*-GFP). In a first set of experiments, human neutrophils were incubated with PTX3-opsonized *S. pneumoniae*-GFP. We confirmed that PTX3 exerts opsonic effects, increasing the phagocytosis of pneumococcus by neutrophils, but only at very high concentrations, i.e. higher than 100 µg/ml (Figure 5B). We then moved to an *in vivo* setting. Since the instillation of as low as 1µg of PTX3 was sufficient to induce an antibacterial effect when administrated locally just before the infection (Figure 4E), we incubated 5×10^4^ CFU of *S. pneumoniae* serotype 3 (i.e. the inoculum normally used for a lethal infection in our model) with 33.3 µg/ml of recombinant PTX3. Mice infected with PTX3-opsonized *S. pneumoniae* serotype 3 showed the same local bacterial burden at 6h to 36h after infection as mice infected with pneumococcus incubated with PBS (Figure 5C). We then evaluated the phagocytic ability of neutrophils recruited *in-vivo* during the infection comparing WT and *Ptx3* deficient mice. Interestingly, we did not observe any difference in the percentage of neutrophils phagocytizing *S. pneumoniae*-GFP in the BAL or in the lung (Figure 5D). Finally, we assessed the killing ability of neutrophils collected from WT and *Ptx3* deficient mice. We did not observe any difference in the percentage of *S. pneumoniae* serotype 3 killed by purified murine neutrophils neither after 1 hour of incubation (WT: 79.02 ± 2.87 and *Ptx3^-/-^*: 80.48 ± 2.87, *P*=0.15) or 3 hours of incubation when nearly all pneumococcus were efficiently killed (WT: 99.46 ± 0.30 and *Ptx3^-/-^*: 96.55 ± 5.40, *P*=0.31) (Figure 5E). These results suggest that the role of PTX3 in resistance to invasive pneumococcus infection is not accounted for by its opsonic activity.

### Regulation of inflammation by PTX3

In pneumococcal invasive disease induced by *S. pneumoniae* serotype 3, infection was characterized by a multifocal neutrophilic bronchopneumonia (Figure S5A). The main inflammatory cell recruitment was observed during the invasive phase of the infection (starting from 24h after infection), when the pulmonary MPO was dramatically increased, (Figure S5B). We analyzed more accurately neutrophil recruitment in the lung and in the BAL of infected mice and we observed two phases of neutrophil recruitment. An initial recruitment, characterized by an increased (i.e. 3-fold compared to uninfected lung) number of neutrophils both in the BAL and in the lung parenchyma, was observed during the first 6h of infection. In the next 12h to 24h of infection we observed an important recruitment of neutrophils in the lung (i.e. 4-fold compared to uninfected lung) that translocated into the alveolar space (up to 50-fold compared to uninfected BAL) (Figure S5C-D). These two steps of recruitment have been described to exert opposite roles (Bou Ghanem et al., 2015). Indeed, the first phase is important for the early control of the infection, reducing the number of colonizing bacteria. In contrast the second phase has been associated with the development of the inflammatory environment, leading to tissue damage that could promote growth and invasion of the bacteria (Bou Ghanem et al., 2015). Given the mild expression of PTX3 during the first hours (Figure 1A), we investigated the second phase of neutrophil recruitment, comparing *Ptx3* deficient and WT mice 18h after infection. At this time point *Ptx3* deficiency was not associated with a higher respiratory bacterial load (Figure S3A). Interestingly, the inflammatory profile was significantly increased in *Ptx3*^-/-^ mice, as shown by an increased development of foci in the lung induced by a higher inflammatory cell recruitment (Figure 6A-C). Moreover, looking at the time course of the development of pneumococcal-induced respiratory inflammation, we observed that *Ptx3*^-/-^ mice had a quicker and more severe formation of inflammatory foci compared to the WT (Figure 6B-C). Furthermore, these mice presented also an increased vascular damage score based on higher perivascular edema and hemorrhages (Figure S5E). Flow cytometry analysis revealed that the higher inflammation in *Ptx3* deficient mice was due to a significant increase of neutrophil recruitment in the BAL and the lung (Figure 6D). Moreover, we did not observe any change in the recruitment of other myeloid cells, i.e. macrophages, eosinophils and monocytes. This phenotype was also observed with the serotype 1 model of invasive pneumococcal infection (Figure S5F).

**Figure 6.**
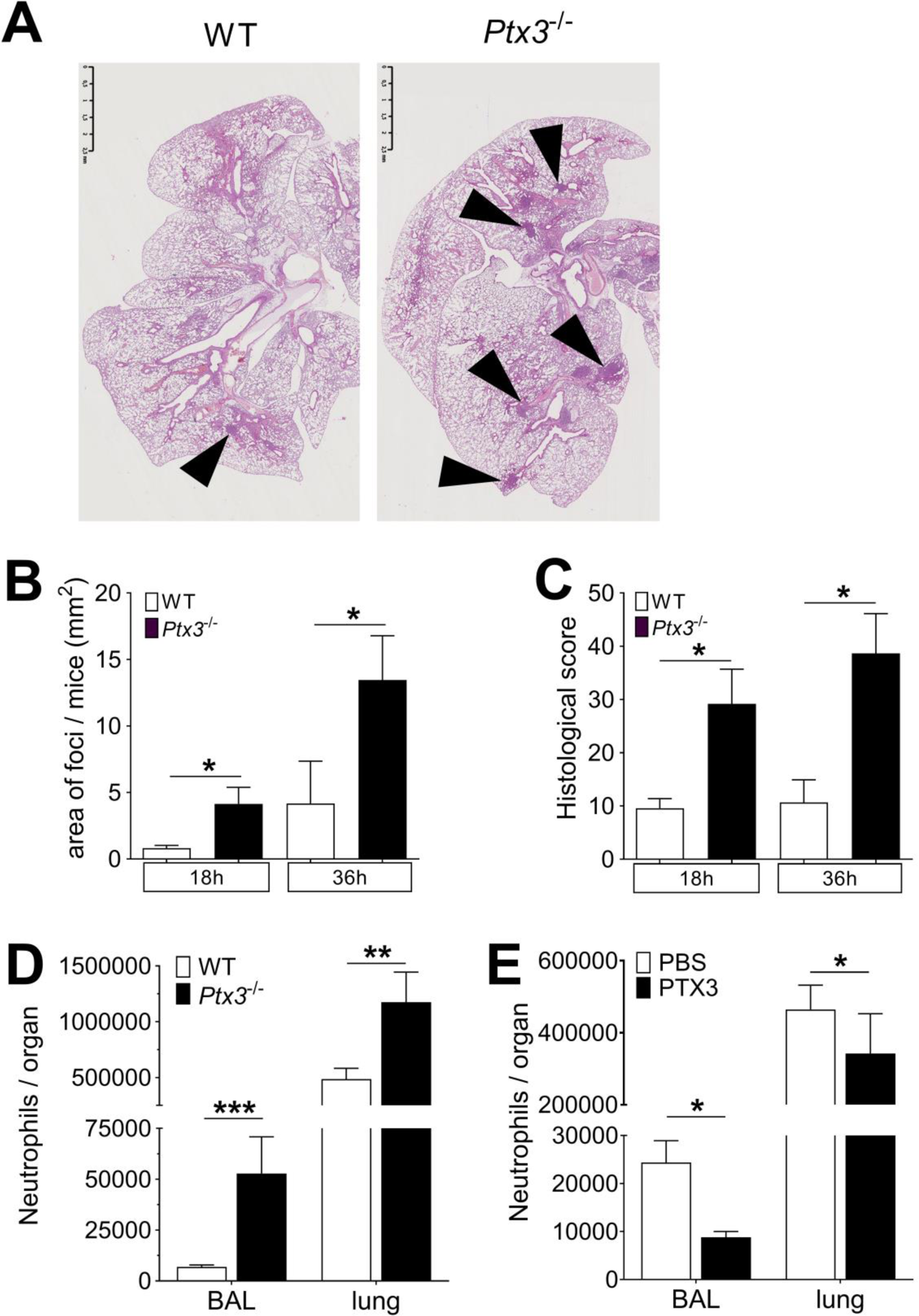
PTX3 regulates inflammation during pneumococcus infection. Mice were infected intranasally with 5×10^4^ CFU of *S. pneumoniae* serotype 3 and sacrificed at the indicated time points for tissue collection. (A) Hematoxylin and Eosin (H&E) staining of formalin-fixed histological sections from the lungs of WT and *Ptx3*^-/-^ mice at 4x magnification. One representative image from at least six biological replicates of WT and *Ptx3^-/-^* mice. Inflammatory cell foci are indicated by arrows. (B) Area of inflammatory cells foci measured in lungs collected 18h and 36h post-infection from WT and *Ptx3*^-/-^ mice. Areas were measured on three H&E stained lung sections per mice at different depth separated by at least 100µm each (n=6-10). (C) Inflammatory histological score measured in lungs collected 18h and 36h post-infection from WT and *Ptx3*^-/-^ mice. Scores (detailed in the Material and Methods section) were determined on three H&E stained lung sections per mice at different depth separated by at least 100µm each (n=6-10). (D) Neutrophil number determined by flow cytometry in BAL and lungs collected 18h post-infection from WT and *Ptx3*^-/-^ mice (data pooled from 2 independent experiments, n=11-18). (E) Neutrophil number determined by flow cytometry in the BAL and lung collected 18h post-infection from WT mice treated intranasally 12h post-infection with recombinant PTX3 or PBS (data pooled from 2 independent experiments, n=11-18). Results represent the mean ± SEM. Statistical significance was determined using the Mann-Whitney test comparing results to uninfected mice (\**P*<0.05, and \*\*\**P*<0.001).

Finally, we observed that intranasal treatment with recombinant PTX3 was also associated with a decrease in the neutrophil number in BAL and lungs, demonstrating that PTX3 has a direct role in the control of neutrophil migration in the respiratory tract (Figure 6E).

### Regulation of neutrophil recruitment by PTX3 during pneumococcal invasive infection

It has been shown that neutrophil depletion during the invasive phase resulted in protection against pneumococcal infection (Bou Ghanem et al., 2015). Accordingly, neutrophil depletion by anti-Ly6G was used to assess the role of these cells in PTX3-mediated protection against pneumococcal infection. In WT mice infected intranasally with *S. pneumoniae,* treatment with anti-Ly6G significantly reduced neutrophils infiltration in the lungs (Figure S6A). In addition, treatment with anti-Ly6G completely abolished the increased accumulation of neutrophils observed in *Ptx3^-/-^* mice (Figure S6A-B). The reduction of neutrophil recruitment in both WT and *Ptx3^-/-^* mice treated with anti-Ly6G resulted in a significant reduction of the local and systemic bacterial load, compared to mice treated with the isotype control (Figure 7A-B). In addition, *Ptx3^-/-^* mice treated with neutrophil depleting antibody were not more infected than the WT mice (Figure 7A-B). These results suggest that taming of pneumococcus-promoting neutrophil recruitment underlies the role of PTX3 in resistance against this bacterial pathogen.

**Figure 7.**
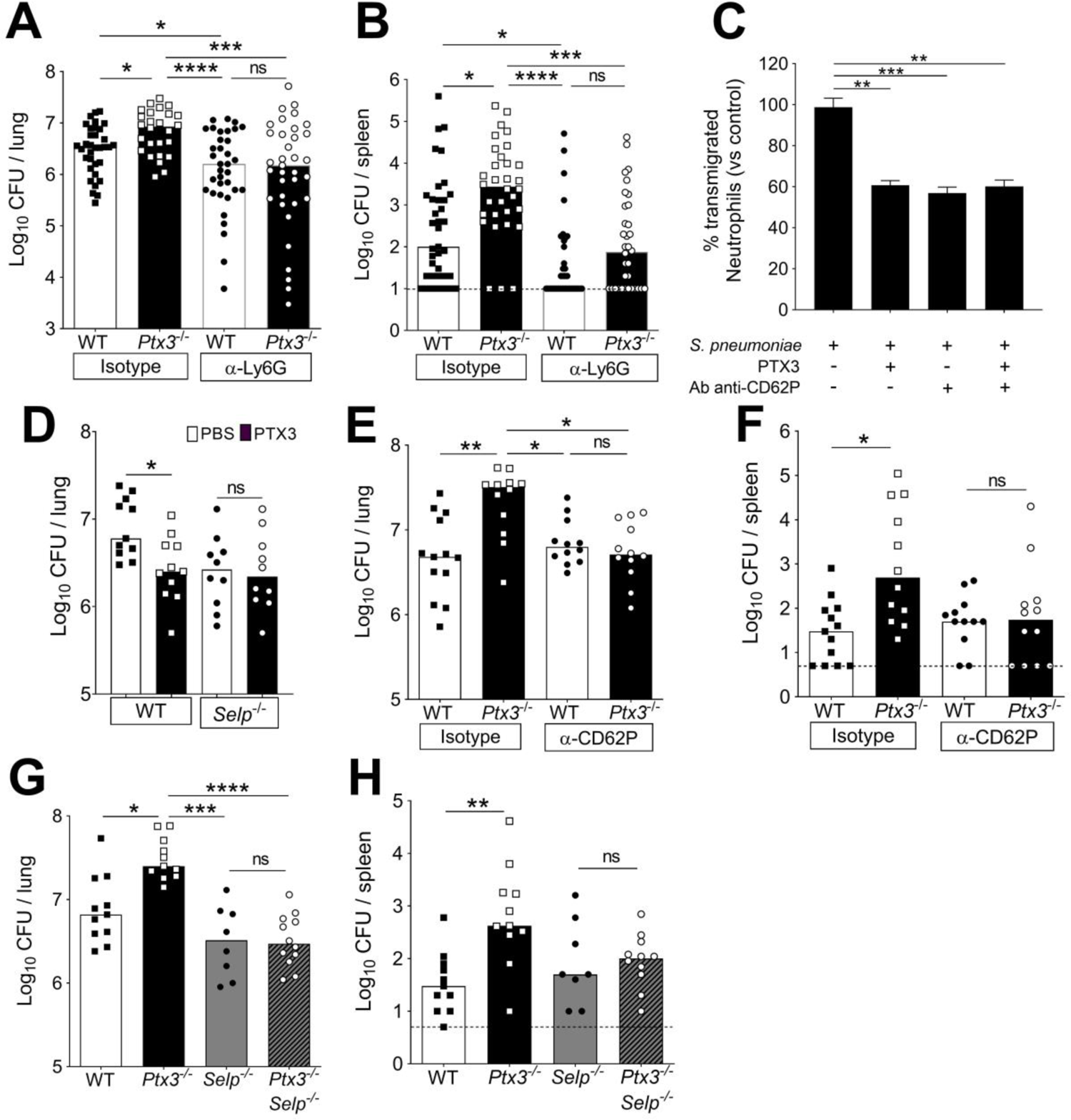
PTX3-mediated regulation of neutrophil recruitment. Mice were infected intranasally with 5×10^4^ CFU of *S. pneumoniae* serotype 3 and sacrificed at the indicated time points for tissue collection. (A-B) Bacterial load in lung (A) and spleen (B) collected 36h post-infection from WT and *Ptx3*^-/-^ mice treated intraperitoneally 12h post-infection with 200µg/100µl of anti-Ly6G or isotype control antibodies (data pooled from 3 independent experiments, n=26-37). (C) Transmigration of human purified neutrophils towards *S. pneumoniae* (data pooled from 2 independent experiments, n=11). Results are reported as percentage of transmigrated neutrophils considering as 100% the number of transmigrated neutrophils in the control condition (i.e *S. pneumoniae* in the lower chamber and no treatment in the upper chamber). (D) Bacterial load in lungs collected 36h post-infection from WT and *Selp*^-/-^ mice treated intranasally 12h post-infection with 1µg/30µl of recombinant PTX3 or PBS (data pooled from 2 independent experiments, n=10-11). (E-F) Bacterial load in lungs (E) and spleens (F) collected 36h post-infection from WT and *Ptx3*^-/-^ mice treated intravenously 12h post-infection with 50µg/100µl of anti-CD62P or isotype control antibodies (data pooled from 2 independent experiments, n=12-13). (G-H) Bacterial load in lungs (G) and spleens (H) collected 36h post-infection from WT, *Ptx3*^-/-^, *Selp*^-/-^ and *Ptx3*^-/-^*Selp*^-/-^ mice (data pooled from 2 independent experiments, n=8-12). Results are reported as median (A-B, D-H) and mean ± SEM. CFU detection limits in the spleen is 5 CFU represented by a dotted line. Statistical significance was determined using the non-parametric Krukal-Wallis test with post-hoc corrected Dunn’s test comparing every means (A-H) (\**P*<0.05, \*\**P*<0.01, \*\*\**P*<0.001 and **** *P*<0.0001).

To dissect the mechanism by which PTX3 orchestrates the modulation of inflammation during pneumococcal infection, we first evaluated the level of neutrophil chemoattractants. At 18h post-infection, even though there was a higher amount of neutrophils in the airways of *Ptx3*^-/-^ mice, we did not detect any differences in the levels of CXCL1 and CXCL2 between *Ptx3* deficient and WT mice (Figure S6C). Since PTX3 is a well-known regulator of complement activation (Haapasalo and Meri, 2019), we investigated the levels of the two anaphylatoxins C3a and C5a in the lung homogenates of infected mice. No difference in the levels of the potent chemoattractants C3a and C5a, was observed (Figure S6C). The levels of C3d, a C3 degradation product deposited on the surface of cells and a marker of complement activation in lung homogenates was similar in *Ptx3* deficient and WT mice (Figure S6D).

PTX3 has been described to directly regulate inflammation by binding P-selectin and reducing neutrophil recruitment, dampening rolling on endothelium (Deban et al., 2010; Lech et al., 2013). We first excluded the presence of any difference in P-selectin levels both in WT and *Ptx3* deficient mice, uninfected or infected with pneumococcus (Figure S6E). Therefore, we investigated whether interaction with P-selectin could be relevant in the regulation of neutrophil recruitment into the lung. We investigated the ability of PTX3 to dampen neutrophil transmigration through endothelial cell layer *in vitro*, using *S. pneumoniae* as the attractive signal. We observed that PTX3 could block 40% of the neutrophil migration induced by *S. pneumoniae* (Figure 7C). Moreover, treatment of endothelial cells with anti-CD62P (P-selectin) antibody induced the same blocking effect. We did not observe any additional blocking effect of PTX3 in association with anti-CD62P, suggesting that PTX3 exerts its blocking effect through P-selectin. To confirm that PTX3 protects infected mice by blocking P-selectin, we used P-selectin deficient mice (*Selp*^-/-^). In *Selp^-/-^* mice PTX3 treatment did not reduce the bacterial load (Figure 7D). Moreover, we treated WT and *Ptx3* deficient mice with anti-CD62P, to block P-selectin-dependent neutrophil transmigration during the invasive phase of infection. Anti-CD62P treatment completely abolished the higher neutrophils recruitment in *Ptx3* deficient mice (Figure S6F-G). This result suggests that the higher neutrophil infiltration observed during pneumococcal pneumonia in the absence of PTX3 is dependent on P-selectin. Importantly, the reduction of neutrophil recruitment in *Ptx3* deficient mice treated with anti-CD62P is associated with a significant reduction of the local and systemic bacterial load reaching the same level observed in WT mice treated with anti-CD62P (Figure 7E-F).

Finally, to assess the role of the P-selectin pathway in PTX3-mediated resistance against invasive pneumococcus infection, we took advantage of *Ptx3*^-/-^*Selp*^-/-^ double deficient mice. As shown in Figure 7G-H, genetic deficiency in P-selectin and PTX3 completely rescued the phenotype observed in *Ptx3^-/-^* mice. Thus, the defective control of invasive pneumococcal infection observed in *Ptx3^-/-^* mice is due to unleashing P-selectin-dependent recruitment of pneumococcus-promoting neutrophils.

### PTX3 polymorphisms

To explore the significance of these results in human, we analyzed the association of human *PTX3* gene polymorphisms with IPD in a cohort of 57 patients and 521 age- and sex-matched healthy controls. We focused in particular on two intronic SNPs (rs2305619 and rs1840680) and a third polymorphism (rs3816527) in the coding region of the protein determining an amino-acid substitution at position 48 (+734A/C). These SNPs are associated with increased susceptibility to infection to selected microorganisms (Chiarini et al., 2010; Cunha et al., 2014; He et al., 2018; Olesen et al., 2007). In addition, the +734A allele was associated in various studies with decreased PTX3 circulating levels (Barbati et al., 2012; Bonacina et al., 2019; Cunha et al., 2014).

Similar frequencies were observed for the +734A allele in patients and controls (67.54% and 61.58%, respectively, *P=*0.213, Table 1). However, when the haplotypes determined by the three SNPs (rs2305619, rs3816527, rs1840680) were examined, we found that the AAA haplotype was twice as frequent in IPD patients as in healthy controls (9.67% and 4.26% respectively, *P*=0.0102, Table 2). This association was even stronger when considering two SNPs only (+281A and +734A), including the one associated with lower levels of the protein (11.4% and 4.94% respectively, *P*=0.0044, Table 2). These observations suggest that also in humans PTX3 could play a role in the control of *S. pneumoniae* infection.

**Table 1.** Frequency distribution of *PTX3* gene single nucleotide polymorphisms (SNPs) in IPD patients and controls.

| SNP | Alleles | Amino acid change | Associated allele | Frequency (%) | | $\chi^2$ | OR (95%) | P-value |
| --- | --- | --- | --- | --- | --- | --- | --- | --- |
|  |  |  |  | IPD patient (n=57) | Control (n=521) |  |  |  |
| rs2305619 | +281 |  | A | 43.86 | 43.44 | 0.007 | 1.02<br>(0.69-1.50) | 0.931 |
|  | A/G |  | G | 56.14 | 56.56 |  |  |  |
| rs3816527 | +734 | Ala→Asp | C | 32.46 | 38.42 | 1.552 | 0.77<br>(0.51-1.16) | 0.213 |
|  | C/A |  | A | 67.54 | 61.58 |  |  |  |
| rs1840680 | +1149 |  | A | 42.11 | 42.49 | 0.006 | 0.98<br>(0.67-1.46) | 0.938 |
|  | A/G |  | G | 57.89 | 57.51 |  |  |  |

**Table 2.**
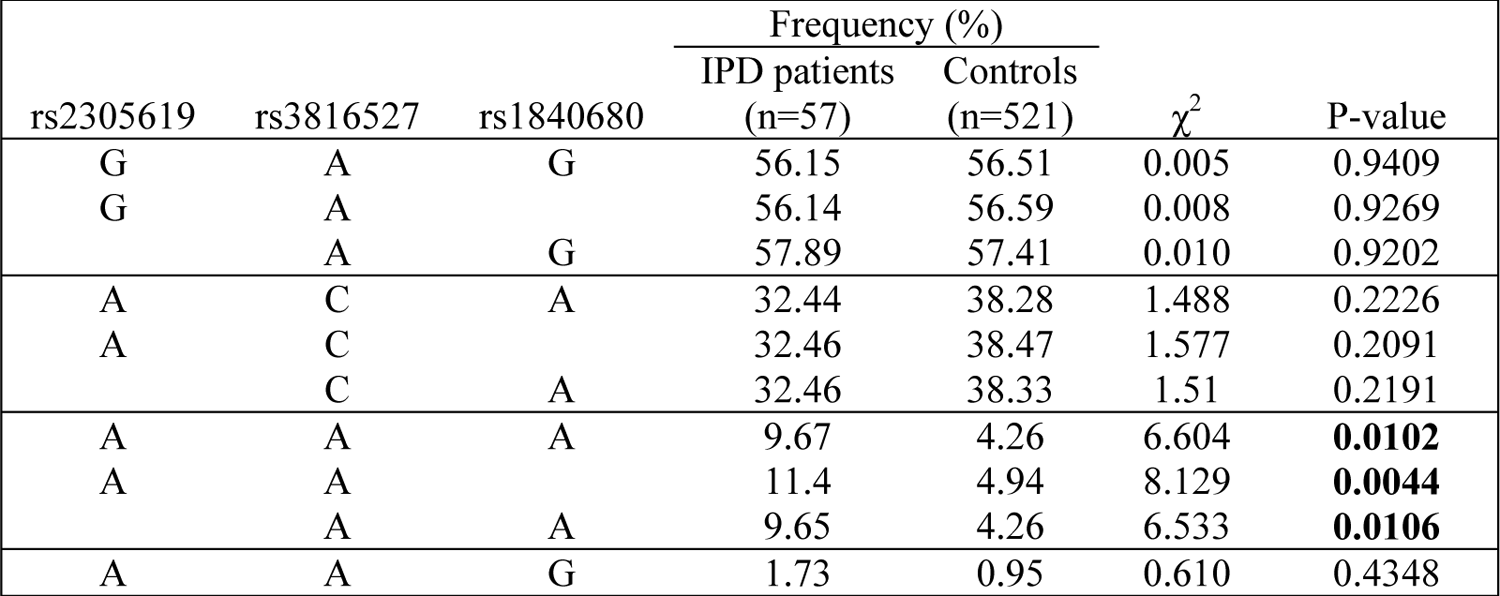
Haplotype analysis for *PTX3* gene in IPD patients and controls.

To assess whether the +734A/C polymorphism in the coding region of the human *PTX3* gene affects the protein’s interaction with P-selectin, two recombinant PTX3 constructs were made that carry either D (Asp) or A (Ala) at position 48 of the preprotein sequence (corresponding to the A and C alleles of the + 734A/C polymorphism, respectively) (Cunha et al., 2014). These two proteins had almost identical electrophoretic mobilities when run on denaturing gels both in reducing and non-reducing conditions (Figure 8A), where they showed a pattern of bands consistent with previous studies (Inforzato et al., 2008). Also, similar chromatograms were recorded when the D48 and A48 variants were resolved on a SEC column in native conditions (Figure 8B). Given that protein glycosylation is a major determinant of the interaction of PTX3 with P-selectin (Deban et al., 2010), it is worth pointing out that the +734A/C polymorphism does not affect structure and composition of the PTX3 oligosaccharides, with major regard to their terminal residues of sialic acid (Bally et al., 2019). Therefore, the allelic variants of the PTX3 protein were virtually identical in terms of quaternary structure and glycosidic moiety, which makes them suitable to comparative functional studies. In this regard, when assayed in solid phase binding experiments, these two proteins equally bound plastic-immobilized P-selectin (Figure 8C), and C1q (Figure 8D, here used as a control), indicating that the +734A/C polymorphism (i.e., the D/A amino acid substitution at position 48 of the PTX3 preprotein) does not affect the interaction of this PRM with P-selectin. It is therefore conceivable that the +734A/C SNP (and the others investigated in our association study) determines reduced expression rather than function of the PTX3 protein *in vivo*, as observed in other opportunistic infections (Cunha et al., 2014).

**Figure 8.**
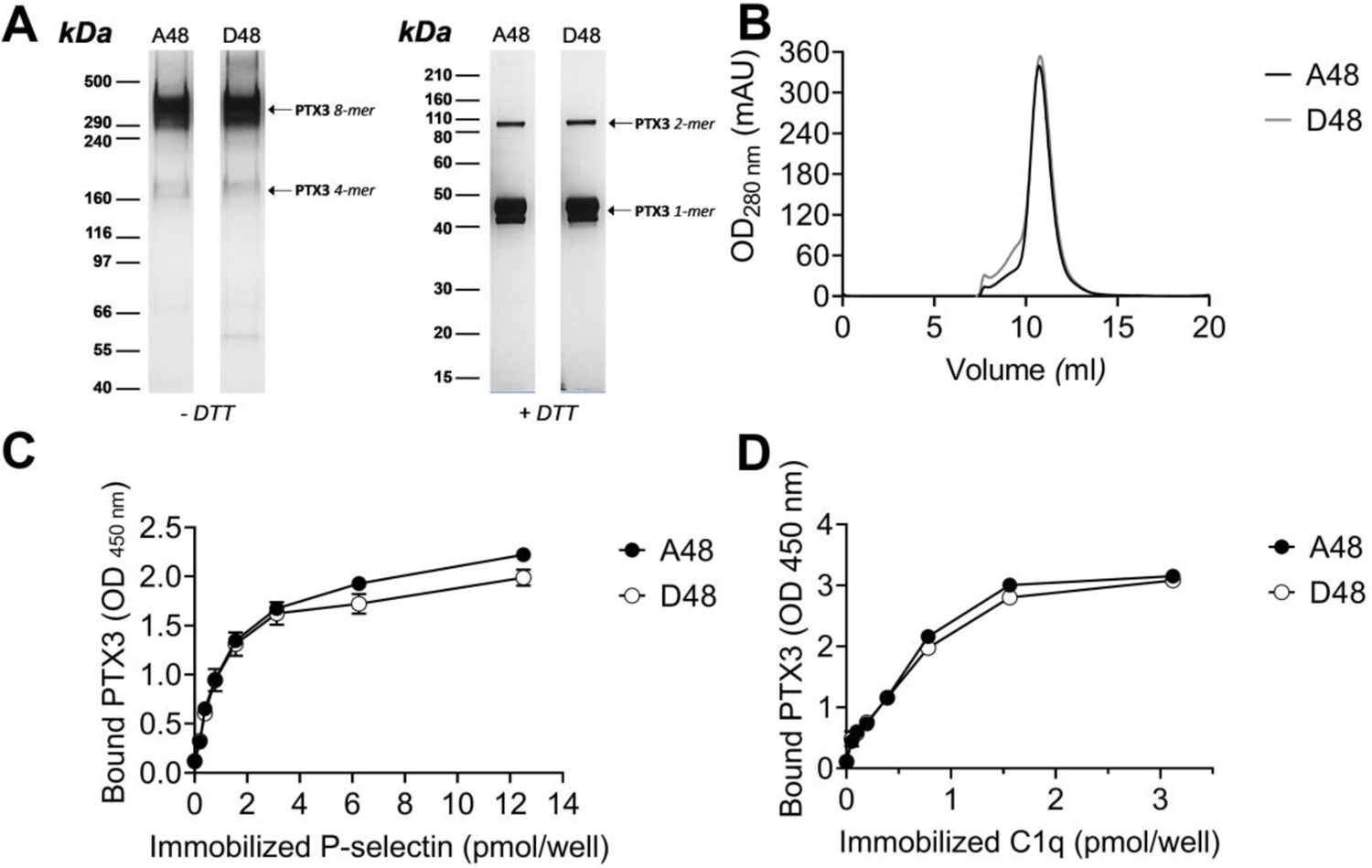
Biochemical characterization of the D48 and A48 allelic variants of PTX3 and their binding to P-selectin. (A) 500 ng/lane of purified recombinant PTX3 (A48 and D48 from HEK293 cells) were run under denaturing conditions on Tris-Acetate 3–8% (w/v) and Bis-Tris 10% (w/v) protein gels, in the absence (-) and presence (+), respectively, of dithiothreitol (DTT). Gels are shown with molecular weight markers on the left, and position of the PTX3 monomers, dimers, tetramers, and octamers (1-, 2-, 4-, and 8-mers, respectively) on the right. (B) 200 µg aliquots of either one of the two allelic variants were separated on a Superose 6 10/300 GL size exclusion chromatography (SEC) column in non-denaturing conditions, with elution monitoring by UV absorbance at 280 nm. An overlay of individual chromatograms is presented. (C and D) The effect of the +734A/C polymorphism on the interaction of PTX3 with P-selectin was investigated by a solid phase binding assay using microtiter plates coated with the indicated amounts of P-selectin or C1q (here, used as a control) that were incubated with either of the A48 and D48 variants (both at 3 nM). Bound proteins were revealed with an anti-human PTX3 polyclonal antibody, and results are expressed as optical density at 450 nm (OD 450 nm), following background subtraction (n=3, mean ±SD). Data shown in panels A to D are representative of three independent experiments with similar results.

## Discussion

*S. pneumoniae* is the most common cause of a range of infections, including community-acquired pneumonia, a pathological condition that affects mainly adults aged 65 years or older and infants under one year of age. It is well known that inflammation plays a crucial role during lung infections and dictates the resolution of pneumonia, but at the same time, exaggerated inflammation can be detrimental (Sohail et al., 2018). Therefore, a strict control of the inflammatory response is essential.

The present study was aimed at assessing the role of PTX3 in invasive pneumococcal infection. PTX3 is a member of the pentraxin family highly conserved in evolution and locally produced by different cell types in response to pro-inflammatory stimuli and microbial components. The protein has multifunctional properties, including in particular a regulatory role on inflammation (Garlanda et al., 2018). In a well-characterized murine model of invasive pneumococcal infection, PTX3 expression was rapidly upregulated in the alveolar and bronchiolar compartments of the lungs. The systemic dissemination of pneumococcus was associated with an increase of PTX3 serum levels. As expected, PTX3 levels were at least in part induced by IL-1β, massively produced in response to pneumococcus.

Both myeloid and endothelial cells can produce PTX3 in response to inflammatory cytokines. Polymorphonuclear leukocytes were able to release PTX3 after stimulation with *S. pneumoniae*, however levels of the protein were similar in WT and in neutropenic *Csfr3^-/-^* animals infected with *S. pneumoniae.* Bone marrow chimeras and conditional mice definitely demonstrated that stromal cells, and in particular endothelial cells, were a major source of PTX3 in this model of pneumococcal infection. Production of PTX3 by non-hematopoietic cells has been previously reported in other experimental settings. In a murine model of arterial thrombosis induced by FeCl_3_, PTX3 was only expressed by vascular cells (Bonacina et al., 2016). Similarly, in a murine model of skin wound-healing, non-hematopoietic cells were the major producers of PTX3 whereas neutrophils showed a minor contribution (Doni et al., 2015).

*Ptx3* genetic deficiency was associated with a higher susceptibility to *S. pneumoniae*. A defective control of bacterial load, associated with a higher mortality rate, was observed during the invasive phase of the infection, and PTX3 administration rescued the phenotype. In humans, *PTX3* gene polymorphisms were already described to have an impact on the susceptibility to selected infections, in particular *Mycobacterium tuberculosis*, *Pseudomonas aeruginosa* and urinary tract infections (Chiarini et al., 2010; Jaillon et al., 2014; Olesen et al., 2007). In addition, *PTX3* genetic variants are associated with the risk to develop invasive aspergillosis or Cytomegalovirus reactivation in patients undergoing allogeneic stem cell transplant (Campos et al., 2019; Cunha et al., 2014). In the present study, in a cohort of 57 patients with IPD and 521 healthy controls haplotypes determined by *PTX3* gene polymorphisms were associated with *S. pneumoniae* infection. Thus, genetic deficiency in mice and genetic polymorphisms in humans suggest that PTX3 plays an important role in the control of invasive pneumococcal infection.

Various mechanisms are potentially involved in the protective role of PTX3 against infectious agents. In most cases PTX3-mediated protection has been related to the pro-phagocytic activity of the protein (Garlanda et al., 2002; Jaillon et al., 2014; Moalli et al., 2011, 2010). PTX3 binds selected fungal, bacterial and viral pathogens, including *Aspergillus fumigatus, Pseudomonas aeruginosa, Shigella flexneri,* uropathogenic *Escherichia coli,* Influenza virus, murine cytomegalovirus as well as SARS-CoV-2 nucleocaspid (Porte et al., 2019; Stravalaci et al., 2022), acting in most cases as an opsonin and amplifying phagocytosis (Garlanda et al., 2002; Jaillon et al., 2014; Moalli et al., 2010). These microorganisms are naturally susceptible to phagocytosis (Hasenberg et al., 2011; Lovewell et al., 2014; Schwab et al., 2017) whereas virulent *S. pneumoniae* developed various mechanisms preventing phagocytosis (Andre et al., 2017; Weiser et al., 2018). In this study, we observed that PTX3 could bind *S. pneumoniae,* promoting its phagocytosis *in vitro* by human neutrophils, only at very high concentrations. *Ptx3* deficiency did not affect the local phagocytosis by recruiting neutrophils and, given the low efficiency of the binding to pneumococcus, pre-opsonisation of the inoculum did not modify the kinetic of infection. Thus PTX3-mediated contribution to resistance to *S. pneumoniae* is independent of enhanced phagocytosis.

The short pentraxin CRP, distantly related to PTX3, acts as an opsonin for various microorganisms, including pneumococcus (Bottazzi et al., 2010; Szalai, 2002). However, this effect is dependent on the serotype, specifically on the expression of phosphatidylcholine in the capsule. PTX3 does not interact with phosphatidylcholine (Bottazzi et al., 1997). In addition, serotypes 1 and 3 gave similar results in terms of kinetic of PTX3 production after infection and bacterial load in *Ptx3^-/-^* mice. These results suggest that the short pentraxin CRP and the long pentraxin PTX3 have distinct spectra of microbial recognition and role in antimicrobial resistance.

PTX3 exerts regulatory roles on complement activation by interacting with components of all the three pathways, i.e. the classical, alternative and lectin pathways. In all cases, PTX3 leads to a reduced activation of the complement cascade, thereby reducing the tissue damage associated with an activation out of control (Haapasalo and Meri, 2019). The higher susceptibility to pneumococcus infection observed in *Ptx3*-deficient mice was not related to failed regulation of complement activity. In fact, similar levels of complement fragments, in particular of the two anaphylatoxins C3a and C5a, were found in lung homogenates of wild type and *Ptx3^-/-^* infected mice.

In invasive pneumococcal infection neutrophils represent a double edged sword. Several lines of evidence, including depletion using anti-Ly6G antibody, suggest that in early phases of infection neutrophils are an essential component of resistance to *S. pneumoniae* as expected (Bou Ghanem et al., 2015). In contrast, during the invasive phase neutrophils depletion was protective, limiting tissue damage and associated bacterial invasion. In the present study, *Ptx3* genetic deficiency was associated with uncontrolled inflammation and bacterial invasion sustained by enhanced neutrophils accumulation and vascular damages, that could lead to an access for pneumococcus to nutrients in the alveolar space allowing pneumococcal outgrowth and dissemination, as already described (Sender et al., 2020). Other studies have shown a protective effect against pneumococcus respiratory infection by controlling lung damage, reducing the neutrophil accumulation and inflammation (Madouri et al., 2018; Porte et al., 2015; Tavares et al., 2016). These results are consistent with a yin/yang role of neutrophils in invasive pneumococcus infection (Nathan, 2006). In contrast with our results, a recent paper reported a proinflammatory role of PTX3 in the context of serotype 2 *S. pneumoniae* infection (Koh et al., 2017). However, when we used D39 serotype 2 pneumococcus in our setting and with our mice, we were not able to find any difference between WT and *Ptx3*^-/-^ pneumococcal induced inflammatory responses, suggesting that other factors, including housing conditions, could have impact on the phenotype.

Neutrophil infiltration at sites of bacterial invasion and inflammation is driven by chemoattractants and adhesion molecules (Maas et al., 2018). Neutrophil attracting chemokines and complement C5a and C3a were no different in *Ptx3^-/-^* and WT mice. PTX3 has been shown to serve as a negative regulator of neutrophil recruitment by interacting with P-selectin (Deban et al., 2010; Lech et al., 2013). *In-vitro* studies and *in-vivo* experiments which took advantage of P-selectin-deficient *Selp^-/-^* mice and *Ptx3*^-/-^*Selp*^-/-^ double deficient mice were designed to assess the relevance of this pathway. The obtained results indicated that the defective control of invasive pneumococcal infection observed in *Ptx3^-/-^* mice is due to unleashing of P-selectin-dependent recruitment of neutrophils which promote bacterial invasion. Thus, by taming uncontrolled P-selectin dependent recruitment of neutrophils, the fluid phase pattern recognition molecule PTX3 plays an essential role in tuning inflammation and resistance against invasive pneumococcus infection.

## Materials and Methods

### Mice

All mice used in this study were on a C57BL/6J genetic background. PTX3-deficient mice were generated as described in (Garlanda et al., 2002). *Ptx3^-/-^* and *P-selectin* (*Selp^-/-^*) double deficient mice were generated as described in (Doni et al., 2015). *Csf3r*^-/-^ mice were generated as described in (Ponzetta et al., 2019). Wild-type (WT) mice were obtained from Charles River Laboratories (Calco, Italy) or were cohoused littermates of the gene-deficient mice used in the study. *Ptx3^-/-^, Csfr3^-/-^, Ptx3loxP^+/+^Cdh5cre^+/+^, Ptx3loxP^+/+^Cdh5cre^-/-^*, *Selp^-/-^, Ptx3^-/-^ Selp^-/-^* and WT mice were bred and housed in individually ventilated cages in the SPF animal facility of Humanitas Clinical and Research Center or purchased from Charles River (Milan) and acclimated in the local animal facility for at least one weeks prior to infection. All animals were handled in a vertical laminar flow cabinet. Procedures involving animals handling and care were conformed to protocols approved by the Humanitas Clinical and Research Center (Rozzano, Milan, Italy) in compliance with national (4D.L. N.116, G.U., suppl. 40, 18-2-1992 and N. 26, G.U. march 4, 2014) and international law and policies (European Economic Community Council Directive 2010/63/EU, OJ L 276/33, 22.09.2010; National Institutes of Health Guide for the Care and Use of Laboratory Animals, U.S. National Research Council, 2011). All efforts were made to minimize the number of animals used and their suffering. The study was approved by the Italian Ministry of Health (742/2016-PR). Experiments were performed using sex- and age-matched mice.

### Bacterial preparation

Each *S. pneumoniae* strain (serotype 1 ST304 and serotype 3 ATCC6303) was cultured and stored as previously described (Porte et al., 2015). Briefly, Todd-Hewitt yeast broth (THYB) (Sigma-Aldrich) was inoculated with fresh colonies grown in blood agar plates and incubated at 37°C until an optical density at 600 nm (OD_600_) of 0.7 to 0.9 units was reached. Cultures were stored at −80°C in THYB with 12% glycerol for up to 3 months. GFP-expressing serotype 1 was constructed as described previously (Kjos et al., 2015). Clinical isolate E1586 serotype 1 *S. pneumoniae* was grown at 37°C in THYE until an OD_600_ of 0.1, then 100 ng/ml of synthetic competence-stimulating peptide 1 (CSP-1; Eurogentec) was added for 12min at 37°C to activate transformation machinery. P*_hlpA_*-*hlpA-gfp*_Cam^r^ DNA fragment provided by Jan-Willem Veening’s group (Kjos et al., 2015) was added to the activated cells and incubated 20min at 30°C. Growth medium was diluted 10 times with fresh THYB medium and incubated 1.5h at 37°C. Transformants were selected by plating 5% sheep blood Tryptic Soy Agar plates (TSA; BD Biosciences) containing 4.5 µg/ml of chloramphenicol, then cultured and stored as described above.

### Mouse model of infection

*S. pneumoniae* serotype 3 and serotype 1 were used to induce pneumococcal invasive infection as described previously (de Porto et al., 2019; Porte et al., 2015). For induction of pneumonia, each mouse was anesthetized by intraperitoneal injection of 100 mg/kg of ketamine plus 10 mg/kg of xylazine in 200µl of PBS. Then 5×10^4^ or 10^6^ colony-forming units (CFU) in 30 µL were inoculated intranasally to induce lethal infection by serotype 3 and serotype 1 respectively. Mouse survival was recorded every 12h. To rescue *Ptx3* deficient mice, they were treated intraperitoneally with 10 µg/200 µl of recombinant PTX3 prior and 24h after infection. Prophylaxis or treatment have been done by intranasal instillation of 1 µg/30 µl recombinant PTX3 prior and 12h after infection respectively. Neutrophil recruitment modulation has been performed by treating intraperitoneally with 200 µg/200 µl of anti-Ly6G depleting antibody (*InVivo*Plus 1A8; BioXcell) or control isotype (*InVivo*Plus rat IgG2a; BioXcell). Blocking of P-selectin was realized by treating intraperitoneally with 50 µg/100 µl of anti-CD62P depleting antibody (rat RB40.34 NA/LE; BD Biosciences) or control isotype (rat IgG1 λ; BD Biosciences).

At indicated time, mice were sacrificed with CO_2_, bronchoalveolar lavage fluid (BAL), serum, lungs, and spleen were harvested and homogenated in PBS for CFU counting or in isotonic buffer (Tris HCl 50 nM, EDTA 2 mM, PMSF 1 mM [Roche Diagnostics GmbH], Triton X-100 1% [Merck Life Science], cOmplete EDTA-free protease inhibitor cocktail [Roche Diagnostics GmbH]) for protein measurement on the supernatant. Bacterial loads per organ were counted by serial dilution plated on 5% sheep blood TSA plates after 12h 37°C 5% CO_2_. Lung CFU were representative of the local infection while splenic CFU were considered as indicator of systemic dissemination of pneumococcus through the bloodstream (Hommes et al., 2014; Porte et al., 2015; Schouten et al., 2014). For histological analysis, the entire lung was collected in organ cassette and fixed overnight in 4% paraformaldehyde (PFA) (immunostaining) or in 10% neutral buffered formalin (hematoxylin eosin staining).

### Recombinant PTX3

Recombinant human PTX3 was purified from culture supernatant of stably transfected Chinese hamster ovary (CHO) cells by immunoaffinity as previously described (Bottazzi et al., 1997). Purity of the recombinant protein was assessed by SDS-PAGE followed by silver staining. Biotinylated PTX3 (bPTX3) was obtained following standard protocols. Recombinant PTX3 contained <0.125 endotoxin units/ml as checked by the Limulus amebocyte lysate assay (BioWhittaker, Inc.). For *in vivo* experiments recombinant PTX3 was diluted in PBS.

To assess the effect on the interaction with P-selectin of the rs3816527 (+734A/C) polymorphism in the human PTX3 gene (that results into a D to A amino acid substitution at position 48 of the preprotein), two constructs were generated by overlapping PCR site-directed mutagenesis, and the corresponding recombinant proteins were expressed in and purified from a HEK293 cell line as previously described (Cunha et al. N Engl J Med. 2014). Aliquots of the purified A48 or D48 PTX3 proteins were run under denaturing conditions on Tris-Acetate 3–8% (w/v) and Bis-Tris 10% (w/v) protein gels (GE Healthcare Life Sciences), in the absence and presence, respectively, of dithiothreitol as reducing agent. Following separation, protein bands were stained with silver nitrate (ProteoSilver Silver Stain Kit, Sigma-Aldrich). The two recombinant proteins were analyzed in non-denaturing conditions on a Superose 6 10/300 GL size exclusion chromatography (SEC) column, equilibrated and eluted with PBS at a flow rate of 0.5 ml/min, using an ÄKTA Purifier FPLC system (GE Healthcare Life Sciences). Protein separation and elution was monitored and recorded by UV absorbance at 280 nm.

### Cell culture and stimulation

Human and murine endothelial cell lines were cultivated to have a confluent monolayer in 12-well culture plates (about 10^5^ cells/well). Human Umbilical Vein Endothelial Cells (HUVEC) were grown in 1% gelatin coated wells in M199 medium (Sigma-Aldrich) containing 20% fetal bovine serum (FBS), 100 µg/ml of Endothelial Cell Growth Supplement (ECGS, Sigma-Aldrich), 100 µg/ml of heparin (Veracer; Medic Italia) and 1% penicillin and streptomycin (Pen/Strep). Murine lung capillary endothelial cell line (1G11) was grown in 1% gelatin coated wells in DMEM with 20% FBS, 100 µg/ml of ECGS, 100 µg/ml of heparin and 1% Pen/Strep.

Human neutrophils were purified from freshly collected peripheral blood in Lithium Heparin Vacutainer (BD Bioscience) and separated by a two-steps gradient separation as previously described by Kremaserova and Nauseef (Quinn and DeLeo, 2020). Briefly leukocytes and erythrocytes were separated by a 3% Dextran from Leuconostoc spp. (Sigma-Aldrich) sedimentation for 40min, then leukocytes in the supernatant were separated with Lympholyte-H Cell Separation Media (Cerdalane) and cells from the lower liquid interphase were rinsed with RPMI.

### Generation of bone marrow chimeras

C57BL/6J wild-type or *Ptx3*-deficient mice were lethally irradiated with a total dose of 900 cGy. Then, 2h later, mice were injected in the retro-orbital plexus with 4×10^6^ nucleated bone marrow cells obtained by flushing of the cavity of a freshly dissected femur from wild-type or *Ptx3*-deficient donors. Recipient mice received gentamycin (0.8 mg/ml in drinking water) starting 10 days before irradiation and maintained for 2 weeks. At 8 weeks after bone marrow transplantation, animals were infected.

### Lung histology and immunostaining

Immunostaining was performed on 8 µm sections from 4% PFA-fixed, dehydrate in sucrose solution and mounted in OCT embedding compound and stored at −80°C. PTX3 staining was performed as described previously (Jaillon et al., 2014). Briefly, sections were stained with 5 µg/ml of rabbit polyclonal antibody anti-human PTX3 as a primary antibody and with MACH 1 universal polymer (Biocare Medical) as a secondary antibody. Staining was revealed with 3,3’Diaminobenzidine (DAB; Biocare Medical) and counterstained with hematoxylin and eosin. Slides were scanned and analyzed with Image-pro (Media Cybernetics) to evaluate the percentage of stained area normalized by analyzing the same area for all animals corresponding to about 25% of the section.

Lung histological analysis was performed on formalin-fixed lungs included in paraffin and 3 µm sections were stained with hematoxylin and eosin. A blind analysis was done on 3 sections per animal distant at least of 150 µm and inflammatory foci were measured determining the area of foci and scores. Scores were determined separating small foci (<0.5 mm) and large foci (>0.5 mm) and then calculating as “Histological score = small foci + large foci x 3”. Vascular damage was scored according to a 5-category scale for perivascular edema and hemorrhage, in which 0 is absent and 1 to 4 correspond to minimal (or focal), mild (or multifocal, <10% of blood vessels), moderate (or multifocal, 10-50% of blood vessels) and marked (or multifocal, >50% of blood vessels), respectively.

### Binding assay

The binding of PTX3 on *S. pneumoniae* was assessed as described previously (Bottazzi et al., 2015). Briefly, 10^6^ CFU *S. pneumoniae* were washed in PBS^+/+^ and suspend with 10 µg/ml, 50 µg/ml or 500 µg/ml of biotinylated recombinant PTX3 for 40min at room temperature. Bacteria were washed with PBS^+/+^ and stained with streptavidin-Alexa Fluor 647 (4 µg/ml, Invitrogen) for 30min at 4°C. Washed bacteria were then fixed with 4% formalin for 15min at 4°C. Bacteria were then read by flow cytometry using FACSCanto II (BD Bioscience). Unstained *S. pneumoniae* were used as negative control.

Binding to P-selectin of the A48 and D48 variants of PTX3 from HEK293 cells was then assessed using 96 well Maxisorp plates (Nunc) coated with a recombinant form of the human P-selectin ectodomain (spanning the 42-771 sequence of the preprotein) commercially available from R&D Systems by adaptation of a published protocol (Bally et al., 2019). Purified C1q from human serum (Merck) was used as a control.

Cells were stimulated after a wash with the same culture media without Pen/Strep and then incubated with the corresponding medium containing 10^6^ CFU *S. pneumoniae*, 20 ng/ml recombinant IL-1β (Preprotech) or 100 ng/ml lipopolysaccharide from *Escherichia coli* O55:B5 (LPS, Sigma-Aldrich) for 6h at 37°C. Cells were then lysate with 300 μl of PureZOL RNA isolation reagent (Bio-Rad). Human neutrophils were stimulated with 10^7^ CFU/ml of *S. pneumoniae* serotype 3 or 10 ng/ml of phorbol myristate acetate (PMA) during 6h at 37°C, PTX3 released in the supernatant was measured by ELISA, as described below.

### Neutrophil transmigration assay

Neutrophil migration assay across an endothelium monolayer was performed as previously described by Bou Ghanem and collaborators (Bou Ghanem et al., 2015). Briefly, basolateral sides of HUVEC monolayer grown 4 days on a 3 µm polyester membrane Transwell (Corning) was infected with *S. pneumoniae* (10^6^ CFU/ml in RPMI) added the lower chamber, whereas 100 µl PBS^+/+^ containing 20 ng/ml recombinant IL-1β supplemented with 100 µg/ml PTX3 and/or 100 µg/ml mouse anti-human CD62P (clone AK-4, BD Bioscience) was added to the apical side (upper chamber). After 2.5h at 37°C, 5×10^5^ human neutrophils (in 100 µl RPMI) were added to the basolateral side. After 2.5h at 37°C, neutrophils in the lower chamber were counted in triplicate. Neutrophil transmigration without infection was performed in parallel as negative control.

### Killing assay

Neutrophil killing of *S. pneumoniae* was evaluated by a resazurin-based cell viability assay using murine purified neutrophils. Briefly, murine neutrophils were purified from bone-marrow as previously descried (Moalli et al., 2010). A volume of 50 µl PBS, containing 4 × 10^5^ CFU *S. pneumoniae* serotype was placed into sterile round bottom Corning 96-well polypropylene microplate and incubated for 1 hour or 3 hours at 37 °C with 100 µl RPMI (10% FBS and GM-CSF 10 ng/ml) containing 2 x 10^5^ murine purified neutrophils from WT and *Ptx3*^-/-^ mice and 50 µl of 10% autologous plasma (WT or *Ptx3*^-/-^) diluted in RPMI +GM-CSF (10 ng/ml). After incubation, plates were immediately cooled on ice and cold-centrifuged, and then supernatant removed. *S. pneumoniae* incubated without neutrophils were used as a negative control. Heat killed (60°C, 2 hours) *S. pneumoniae* were considered as positive control in the assay. Neutrophils were then lysate with 200 µl of distillated water and vigorous shaking. Remaining *S. pneumoniae* were then suspend in 20 µl RPMI. Preparation of AlamarBlue Cell Viability Reagent and test was performed according with manufacturer’s instructions (ThermoFisher Scientific-Invitrogen). A volume of 180 µl AlamarBlue solution (18 µl of AlamarBlue reagent and 162 µl of RPMI) was added to each well. After 4 hour incubation at 37 °C, fluorescence (excitation/emission at ≈530– 560/590 nm) intensity was measured by microplate reader Synergy H4 (BioTek, France). Results represent ratio of fluorescence intensity values relative to those measured in negative controls.

### Gene expression quantification by real-time RT-PCR

Organs homogenated in PureZOL RNA isolation reagent (Bio-Rad) and cell lysate RNAs were extracted with the Direct-zol RNA Miniprep (Zymo Research) and reverse transcribed with the high-capacity cDNA archive kit (Applied Biosystems) following the manufacturer’s instructions. cDNA was amplified using the Fast SYBR Green Master Mix on a QuantStudio 7 Flex Real Time PCR Systems (Applied Biosystems). The sequences of primer pairs (Sigma-Aldrich) specific for murine *Gapdh* (Forward, 5’-GCAAAGTGGAGATTGTTGCCAT-3’, Reverse, 5’-CCTTGACTGTGCCGTTGAATTT-3’) and *Ptx3* (Forward, 5’-CGAAATAGACAATGGACTCCATCC-3’, Reverse, 5’-CAGGCGCACGGCGT-3’) were used to evaluated their expression. Relative mRNA levels (2^-ΔΔCT^) were determined by comparing first the PCR cycle thresholds (CT) for *Ptx3* and *Gapdh* (ΔCT), and second, the ΔCT values for the infected/treated and uninfected/untreated (mock/control) groups (ΔΔCT). All amplifications were performed in triplicates.

### ELISA

Lung homogenates and serum levels of murine C3a, C5a, CXCL1, CXCL2, IL-1β, MPO, PTX3 and P-selectin were determined by enzyme-linked immunosorbent assay (DuoSet ELISA, R&D Systems and Cloud-Clone corp) following the manufacturer’s instructions. Human PTX3 was determined with an in-house ELISA as previously described by Jaillon and collaborators (Jaillon et al., 2014). Briefly, anti-PTX3 monoclonal antibody (1 µg/ml, clone MNB4) in carbonate buffer (carbonate buffer 15 mM pH 9.6) was coated overnight at 4°C in 96 well ELISA plates (Nunc). Wells were then blocked with 5% dry milk for 2h at room temperature. Cell culture supernatants, were incubated for 2h at room temperature. Biotin-labeled polyclonal rabbit anti-PTX3 antibody (100 µg/ml) was used for the detection and incubated 1h at 37°C. Plates were incubated with peroxidase-labeled streptavidin (SB01-61; Biospa) for 1h at 37°C. Bound antibodies were revealed using the TMB substrate (Sigma Aldrich) and 450 nm absorbance values were read with an automatic ELISA reader (VersaMax; Molecular Devices).

### Flow cytometry

BAL fluid samples were obtained after intratracheal injection of 1 ml of PBS supplemented with 5% FBS. Lung cells were isolated after digestion in PBS, supplemented with 20% FBS, 2 mM HEPES (Lonza), 100 µg/ml collagenase from Clostridium histolyticum type IV (Sigma-Aldrich) and 20 µg/ml of DNAse (Roche Diagnostics GmbH) in C-tubes processed with gentleMACS Octo Dissociator with heaters according to the manufacturer’s instructions (Miltenyi Biotec). Lysate were pellet (500 g 8min) and red blood cells were lysate with 500 µl of ACK lysing buffer (Lonza) for 5min. Reaction were stopped with PBS, the cell suspensions were filtered through a 70 μm filter, count using Türk solution (Sigma-Aldrich) and 10^6^ cells were pelleted by centrifugation (500 g, 8 min). Live/dead fixable aqua (Invitrogen) staining were realized following manufacturer’s instruction and stopped in FACS buffer (PBS, 2% FBS, 2 mM EDTA, 0.05% NaN_3_). Fc-receptors were blocked with anti-mouse CD16/CD32 (20 µg/ml, clone 93; Invitrogen) for 20min. Cells were stained with an antibody panel able to distinguish macrophages (CD45^+^, CD11b^-^, SiglecF^+^), neutrophils (CD45^+^, CD11b^+^, SiglecF^-^, Ly6C^+^, Ly6G^+^), monocytes (CD45^+^, CD11b^+^, SiglecF^-^, Ly6C^low/moderate/high^, Ly6G^-^) and eosinophils (CD45^+^, CD11b^+^, SiglecF^+^) as described in Figure S7: anti-CD45-Brilliant Violet 605 (2 µg/ml, clone 30-F11; BD Bioscience), anti-CD11b APC-Cy7 (1 µg/ml, clone M1/70; BD Bioscience), anti-SiglecF-eFluor 660 (1.2 µg/ml, clone 1RNM44N; Invitrogen), anti-Ly6C-FITC (3 µg/ml, clone AL-21; BD Bioscience), anti-Ly6G-PE-CF594 (0.4 µg/ml, clone 1A8; BD Bioscience). Flow cytometric analysis was performed on BD LSR Fortessa and analyzed with the BD FACSDiva software.

### Genotyping

DNA was obtained from 57 pediatric patients with invasive pulmonary disease (IPD) and 521 age- and sex-matched healthy controls from the cohort described by Garcia-Laorden and collaborators (García-Laorden et al., 2020). The genotyping was performed as previously described by Barbati and collaborators (Barbati et al., 2012). Briefly, genomic DNAs extracted from frozen EDTA-whole blood were genotyped by real time-PCR, using TaqMan. In particular, 5 µl samples containing TaqMan Genotyping Master Mix, and specific TaqMan SNP genotyping probes (rs1840680, rs2305619 and rs3816527) were mixed with 20 ng of genomic DNA and genotyped using a Quantstudio 6 Flex System according to the manufacturer’s instruction (Applied Biosystems).

### Statistical analysis

Results were expressed as median or mean ± SEM as indicated. Statistical differences were analyzed using the non-parametric Mann-Whitney test for two groups comparison, or the non-parametric Krukal-Wallis test with post-hoc corrected Dunn’s test for multiple comparison of the mean with unequal sample size; survival analysis was performed with the logrank test with Mantel-Cox method. All the analyses were performed with GraphPad Prism 8.0; *P* values <0.05 were considered significant.

Sample size estimation was determined for each read-out by performing pilot experiments and determining the Cohen’s effect size *d* (Lakens, 2013). Sample size were then estimated using G*Power software (version 3.1.9.7) to perform an *a priori* power analyses considering the *d* calculated as described above, an α error probability of 0.05 and 0.01 and a power level (1-β error probability) of 0.8 and considering the appropriated statistical analyses test (Faul et al., 2007). Depending on the model, the sample size ranges between 3 and 40. Number of animals used are reported in the appropriate legends to figures.

As for SNP association analyses, these were performed using the PLINK v1.07 program (Purcell et al., 2007). All polymorphisms had a call rate of 100%, and were tested for Hardy-Weinberg equilibrium (HWE) in controls before inclusion in the analyses (*P*-HWE >0.05). In detail, deviations from HWE were tested using the exact test (Wigginton et al., 2005) implemented in the PLINK software. For each SNP, a standard case-control analysis using allelic chi-square test was used to provide asymptotic *P* values, odds ratio (OR), and 95% confidence interval (CI), always referring to the minor allele. Haplotype analysis and phasing was performed considering either all three SNPs together or by using the sliding-window option offered by PLINK. All *P* values are presented as not corrected; however, in the relevant tables, Bonferroni-corrected thresholds for significance are indicated in the footnote.

## Acknowledgements

The financial support of Fondazione Cariplo (Contract n° 2015-0564), the Italian Spacial Agency (ASI - MARS-PRE Project, grant number DC-VUM-2017-006), and Associazione Italiana Ricerca sul Cancro (AIRC –grant IG-2019 Contract n° 23465) are gratefully acknowledged. We also acknowledge Jean-Claude Sirard team “Bacteria Antibiotics and Immunity”, Center for Infection and Immunity of Lille, France, for providing serotype 1 pneumococcal strain, and Tom van der Poll team, Academic Medical Center of Amsterdam, Netherlands, for providing us serotype 3 pneumococcal strain. CG, FA, BB and AM are supported by the European Sepsis Academy Horizon 2020 Marie Skłodowska-Curie Action: Innovative Training Network (MSCA-ESA-ITN, grant number 676129). A.R.G. received financial support from Fundação para a Ciência e a Tecnologia (FCT) for PhD grants PD/BD/114138/2016. CT received a scholarship from the Société Académique Vaudoise (Lausanne, Switzerland). The financial support of Fondazione Beppe e Nuccy Angiolini to RaPa and AI is greatly acknowledged.

## Supplementary Figures

**Figure S1.**
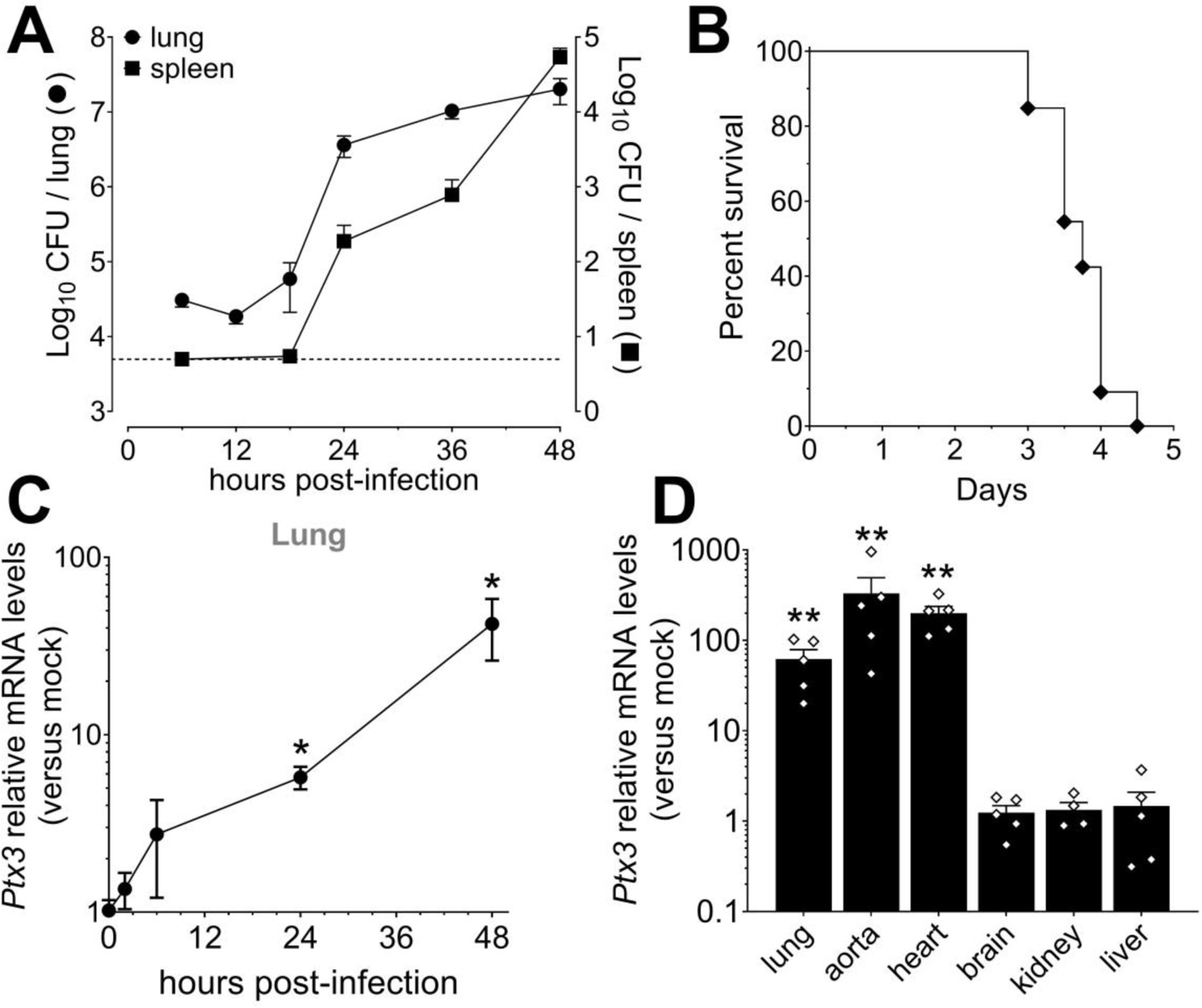
Invasive pneumococcal infection induces PTX3 expression. Mice were infected intranasally with 5×10^4^ CFU of *S. pneumoniae* serotype 3 and sacrificed at the indicated time points for tissue collection. (A) Bacterial load in lung (●) and spleen (▪) collected at indicated time points after infection of WT mice (n=10-21). (B) Survival of WT mice infected with a lethal inoculum of *S. pneumoniae* serotype 3. Mice were monitored every 6h (n=33). (C) Relative *Ptx3* mRNA expression determined by Real-Time quantitative PCR in lung homogenates collected at the indicated time points and normalized on uninfected mice (n=3). (D) Relative *Ptx3* mRNA expression determined by Real-Time quantitative PCR in the indicated organ homogenates collected 48h post-infection and compared to uninfected mice mice (n=4-5). Results are reported as the mean ± SEM. CFU detection limits in the spleen is 5 CFU represented by a dotted line. Statistical significance was determined using the Mann-Whitney test comparing results to uninfected mice (\**P*<0.05 and \*\**P*<0.01).

**Figure S2.**
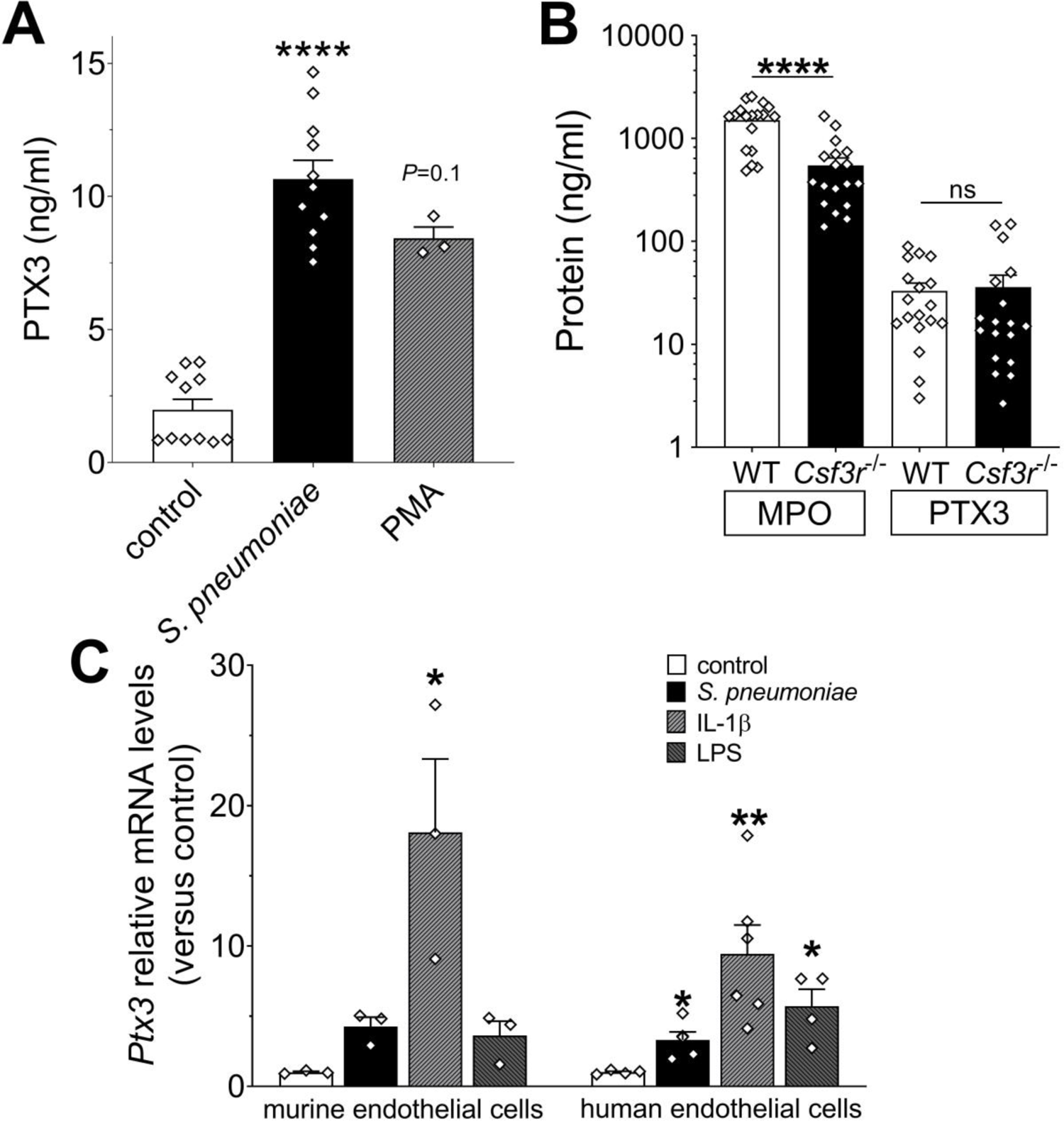
Cellular sources of PTX3 after stimulation with *S. pneumoniae*. (A) PTX3 protein levels, measured by ELISA, released by 10^6^ human purified neutrophils/100µl stimulated for 6h at 37°C with 10^7^ CFU/ml of *S. pneumoniae* serotype 3 or 10 ng/ml of phorbol myristate acetate (PMA). (B) MPO and PTX3 protein levels determined by ELISA in lung homogenates collected 36h post intranasal infection of WT and *Csfr3*^-/-^ mice with 5×10^4^ CFU of *S. pneumoniae* serotype 3 (data pooled from 2 experiments, n=18). (C) Relative *Ptx3* mRNA expression determined by Real-Time quantitative PCR in human and murine endothelial cells after 6h stimulation with 10^6^ CFU *S. pneumoniae*, 20 ng/ml IL-1β or 100 ng/ml LPS (n=3-6). Results are reported as mean ± SEM. Statistical significance was determined using the non-parametric Krukal-Wallis test with post-hoc corrected Dunn’s test comparing means to control group (A, C) or the Mann-Whitney test (B) (\**P*<0.05, \*\**P*<0.01 and **** *P*<0.0001).

**Figure S3.**
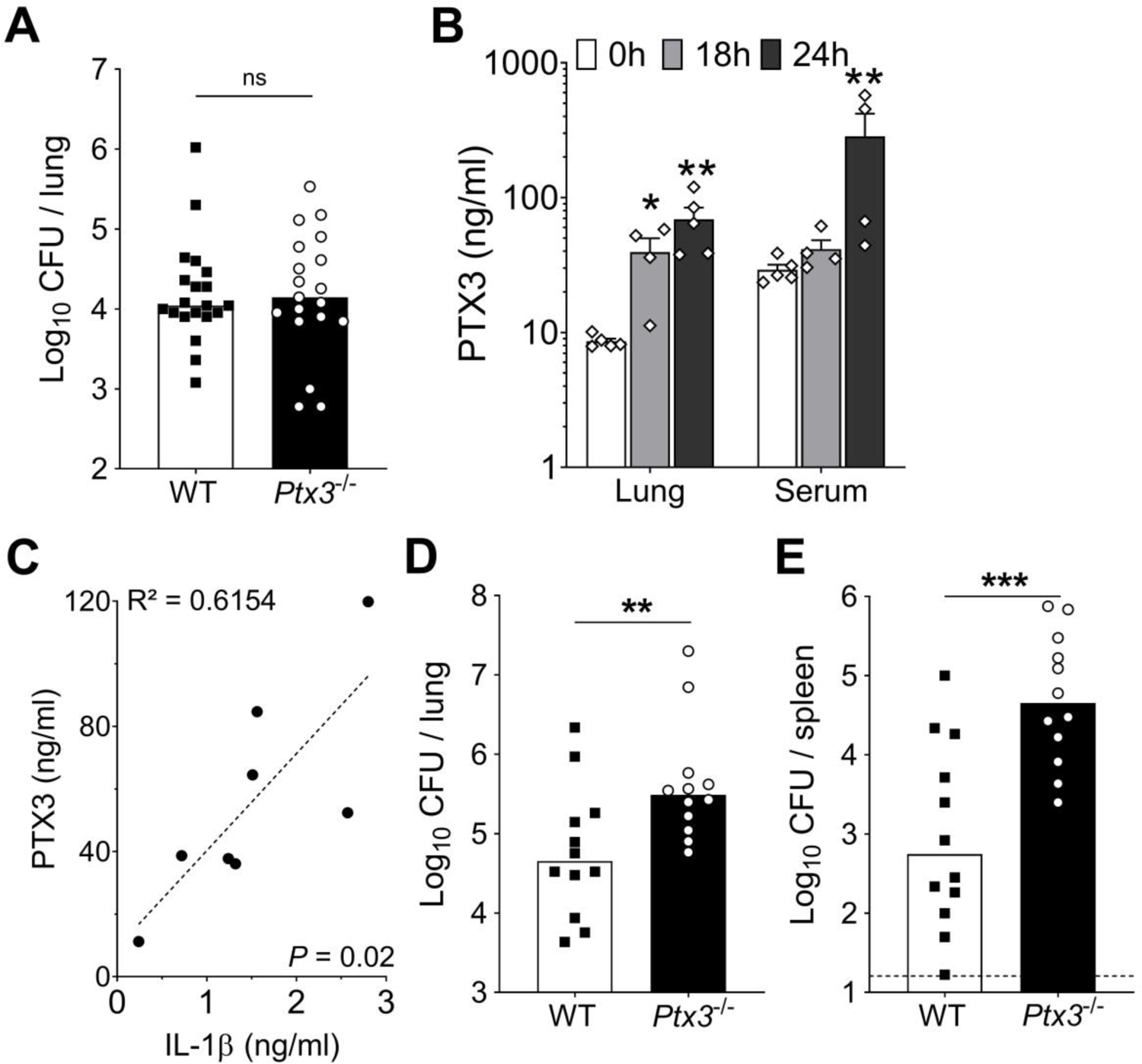
Infection with *S. pneumoniae* serotype 1 and bacterial growth rate in the presence of PTX3. Mice were infected intranasally with 5×10^4^ CFU of *S. pneumoniae* serotype 3 (A) or with 10^6^ CFU of *S. pneumoniae* serotype 1 (B-E) and sacrificed at the indicated time points for tissue collection. (A) WT and *Ptx3*^-/-^ mice were infected with 5×10^4^ CFU and bacterial load in lung was analyzed at 18h post-infection (data pooled from 2 independent experiments, n=19-20). (B) PTX3 protein levels determined by ELISA in lung homogenates and serum collected at the indicated time points (n=4-5). (C) Correlation between PTX3 and IL-1β protein levels in lung homogenates of all infected mice sacrificed from 18 to 24 hours post-infection (n=8). (C-D) Bacterial load in lung (D) and spleen (E) collected at the indicated time points after infection of WT and *Ptx3*^-/-^ mice with *S. pneumoniae* serotype 1 (n=12). Results are reported as the mean ± SEM (B). CFU detection limits in the spleen is 5 CFU represented by a dotted line. Statistical significance was determined using the non-parametric Krukal-Wallis test with post-hoc corrected Dunn’s test comparing means to uninfected mice (B) and the Mann-Whitney test (A, D-E) (\**P*<0.05, \*\**P*<0.01 and \*\*\**P*<0.001).

**Figure S4.**
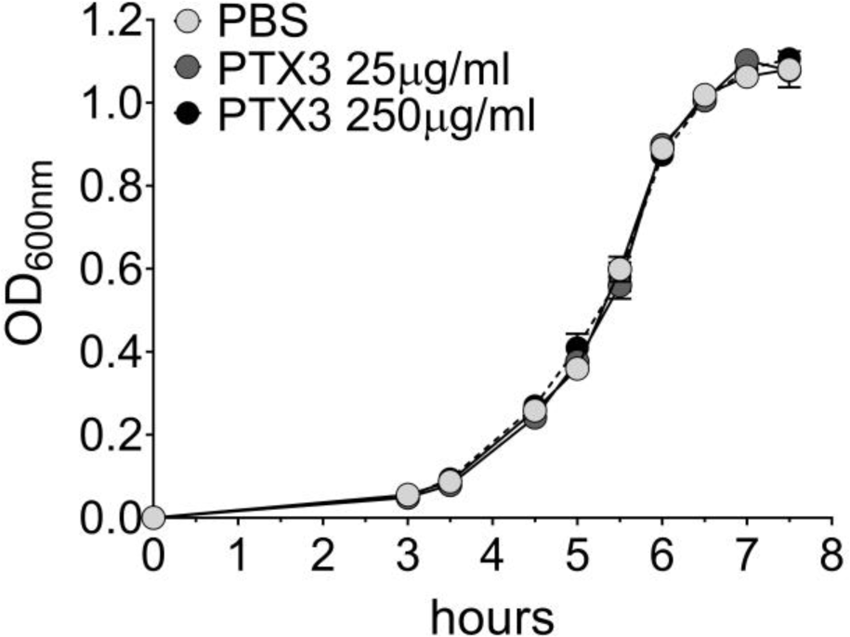
Lack of effect of PTX3 on *S. pneumoniae* growth rate. Growth rate of *S. pneumoniae* serotype 3 non-opsonized or pre-opsonized with recombinant PTX3 (25-250 µg/ml for 40 min) was measured in the culture condition reported in the Material & Methods section. Absorbance (600 nm) was measured at the indicated time points (n=3) and is reported as mean ± SEM.

**Figure S5.**
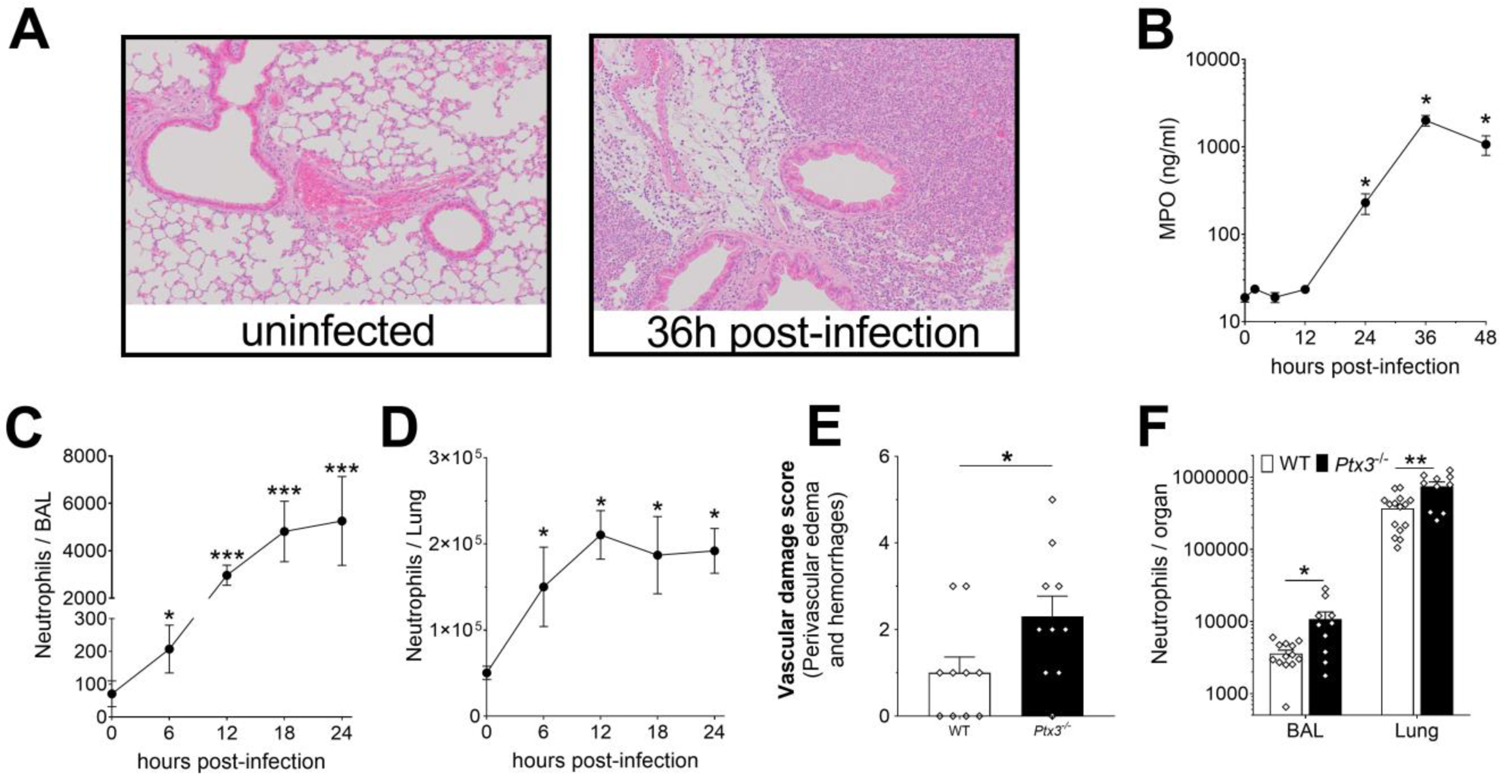
Neutrophil recruitment during invasive pneumococcus infection. Mice were infected intranasally with 5×10^4^ CFU of *S. pneumoniae* serotype 3 (A-E) or with 10^6^ CFU of serotype 1 (F) and sacrificed at the indicated time points for tissue collection. (A) Hematoxylin and Eosin (H&E) staining of formalin-fixed lung sections from WT mice uninfected and 36h after infection at 10x magnification. (B) MPO levels determined by ELISA in lung homogenates collected at the indicated time points (n=4). (C-D) Neutrophil number determined by flow cytometry in the BAL (C) and lung (D) collected at the indicated time points from WT mice (n=4-8). (E) Vascular damage histological score measured in lungs collected 18h post-infection from WT and *Ptx3*^-/-^ mice. Scores (detailed in the Material and Methods section) were determined on three H&E stained lung sections per mice at different depth separated by at least 100µm each (n=6-10). (F) Neutrophil number determined by flow cytometry in the BAL and lung collected 18h post-infection from WT and *Ptx3*^-/-^ mice (data pooled from 2 independent experiments, n=10-13). Results are reported as mean ± SEM. Statistical significance was determined using the Mann-Whitney test comparing results to uninfected mice (\**P*<0.05, \*\**P*<0.01 and \*\*\**P*<0.001).

**Figure S6.**
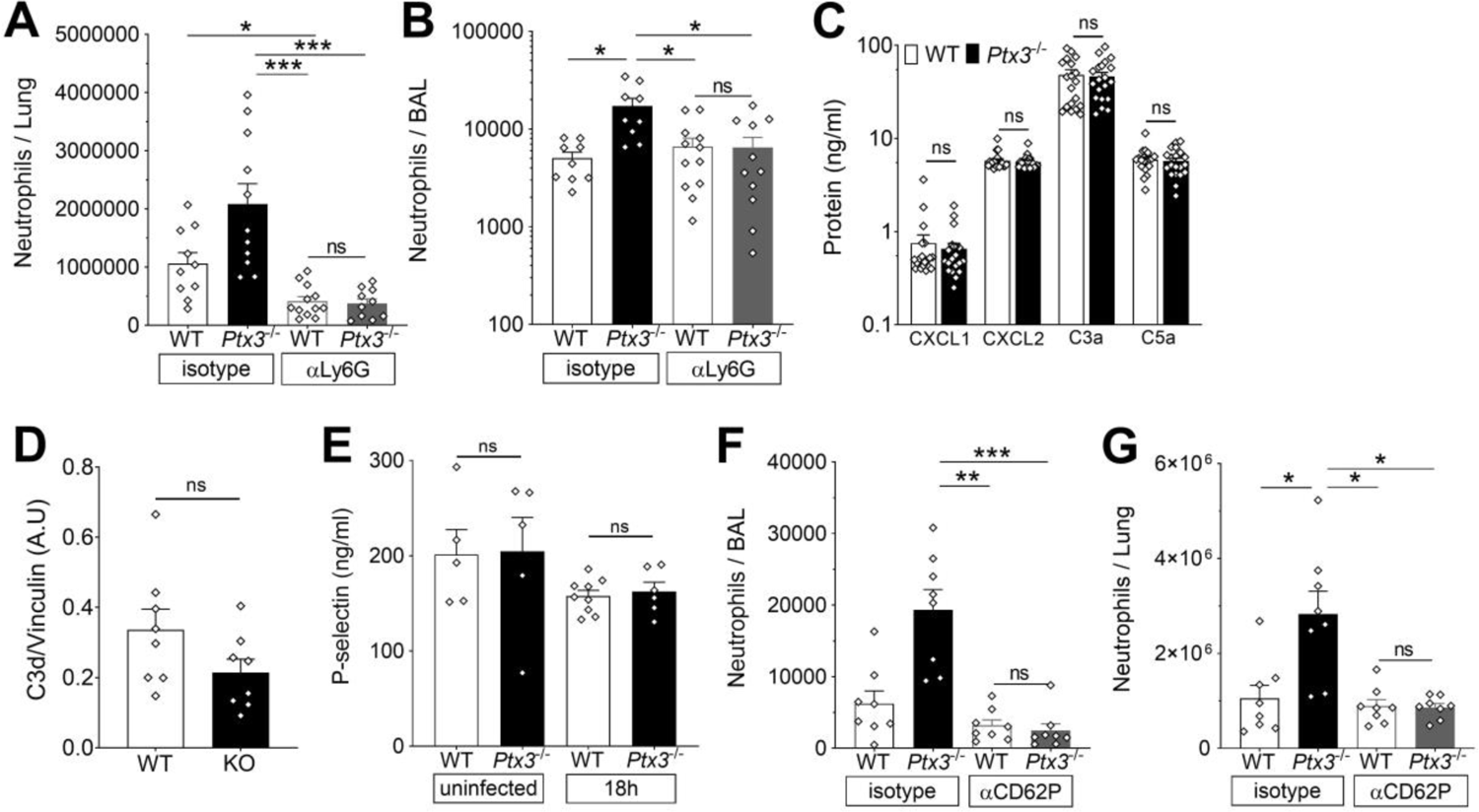
PTX3 modulates neutrophil recruitment. Mice were infected intranasally with 5×10^4^ CFU of *S. pneumoniae* serotype 3 and sacrificed at the indicated time points for tissue collection. (A-B) Neutrophil number determined by flow cytometry in the lung (A) and BAL (B) collected 18h post-infection from WT and *Ptx3*^-/-^ mice treated intraperitoneally 12h post-infection with 200µg/100µl of anti-Ly6G or isotype control antibodies (data pooled from 2 independent experiments, n=9-12). (C) Chemokines (CXCL1/CXCL2) and anaphylatoxins (C3a/C5a) levels measured by ELISA in lung homogenates collected 18h post-infection from WT and *Ptx3*^-/-^ mice (data pooled from 2 independent experiments, n=20). (D) C3d level in lung homogenates collected 36h post-infection from WT and *Ptx3*^-/-^ (KO) mice, detected by western blot and normalized with vinculin expression (n=8). (E) P-selectin expression in lung at steady state or during *S. pneumoniae* respiratory infection. WT and *Ptx3^-/-^* mice were infected intranasally with 5×10^4^ CFU of S*. pneumoniae* serotype 3 and sacrificed 18h post-infection for lung tissue collection. Uninfected mice were also collected for steady state expression. P-selectin expression was evaluated in lung homogenates by ELISA (n=5-9). (F-G) Neutrophil number determined by flow cytometry in the BAL (F) and lung (G) collected 18h post-infection from WT and *Ptx3*^-/-^ mice treated intraperitoneally 12h post-infection with 50µg/100µl of anti-CD62P or isotype control antibodies (n=8). Results are reported as mean ± SEM. Statistical significance was determined using the non-parametric Krukal-Wallis test with post-hoc corrected Dunn’s test comparing means to WT mice treated with isotype antibody (A-B, E-G) and the Mann-Whitney test (C) (\**P*<0.05, \*\**P*<0.01 and \*\*\**P*<0.001).

**Figure S7.**
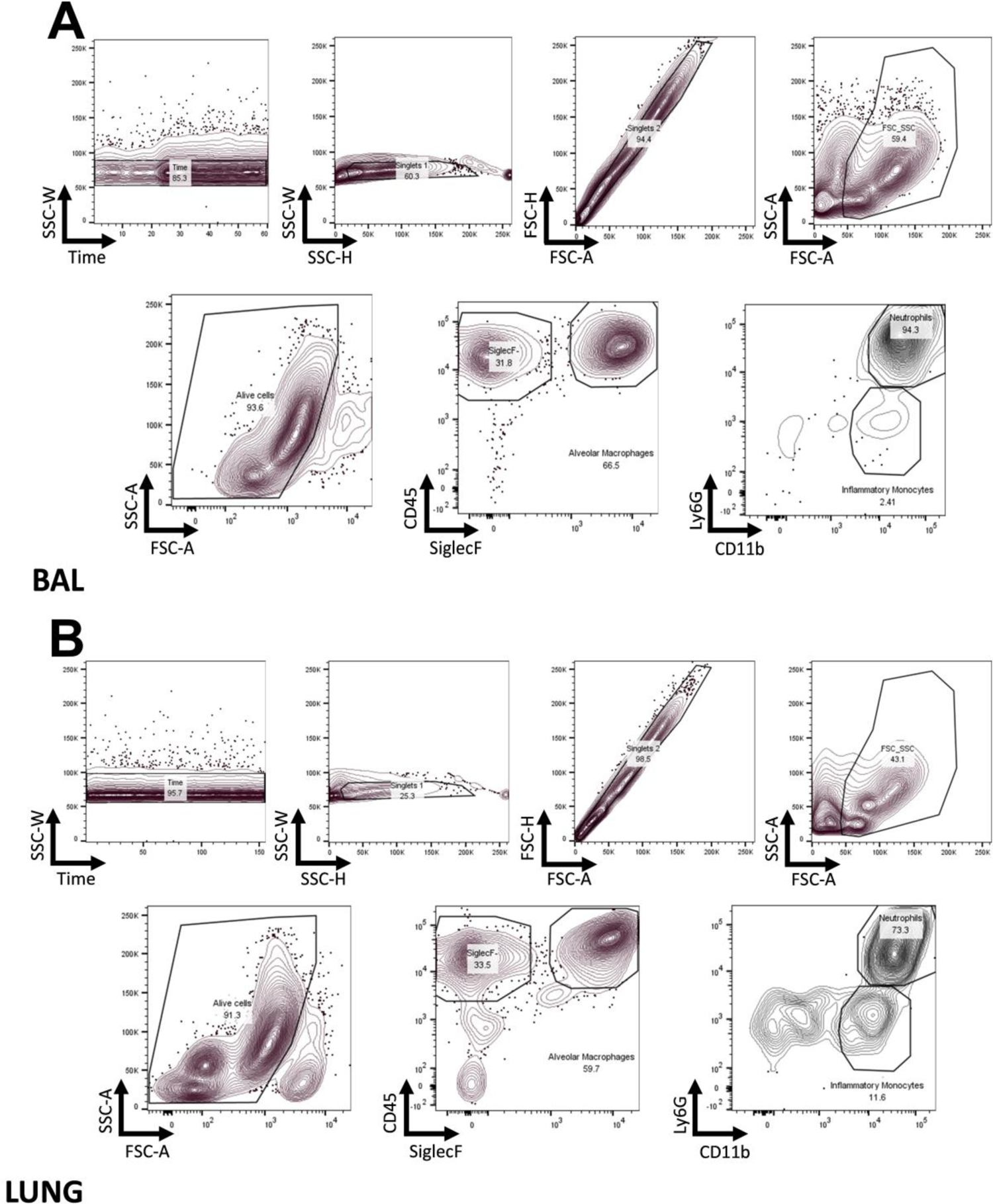
FACS gating strategy. Identification of myeloid subset in the BAL and in the lung reported in the Material & Methods section.

## References

1. Andre GO, Converso TR, Politano WR, Ferraz LFC, Ribeiro ML, Leite LCC, Darrieux M. 2017. Role of Streptococcus pneumoniae Proteins in Evasion of Complement-Mediated Immunity. Front Microbiol 8. doi:10.3389/fmicb.2017.00224

2. Bally I, Inforzato A, Dalonneau F, Stravalaci M, Bottazzi B, Gaboriaud C, Thielens NM. 2019. Interaction of C1q With Pentraxin 3 and IgM Revisited: Mutational Studies With Recombinant C1q Variants. Front Immunol 10:461. doi:10.3389/fimmu.2019.00461

3. Barbati E, Specchia C, Villella M, Rossi ML, Barlera S, Bottazzi B, Crociati L, d’Arienzo C, Fanelli R, Garlanda C, Gori F, Mango R, Mantovani A, Merla G, Nicolis EB, Pietri S, Presbitero P, Sudo Y, Villella A, Franzosi MG. 2012. Influence of Pentraxin 3 (PTX3) Genetic Variants on Myocardial Infarction Risk and PTX3 Plasma Levels. PLoS ONE 7:e53030. doi:10.1371/journal.pone.0053030

4. Bilgin H, Haliloglu M, Yaman A, Ay P, Bilgili B, Arslantas MK, Ture Ozdemir F, Haklar G, Cinel I, Mulazimoglu L. 2018. Sequential Measurements of Pentraxin 3 Serum Levels in Patients with Ventilator-Associated Pneumonia: A Nested Case-Control Study. Canadian Journal of Infectious Diseases and Medical Microbiology 2018:1–8. doi:10.1155/2018/4074169

5. Bonacina F, Barbieri SS, Cutuli L, Amadio P, Doni A, Sironi M, Tartari S, Mantovani A, Bottazzi B, Garlanda C, Tremoli E, Catapano AL, Norata GD. 2016. Vascular pentraxin 3 controls arterial thrombosis by targeting collagen and fibrinogen induced platelets aggregation. Biochimica et Biophysica Acta (BBA) - Molecular Basis of Disease 1862:1182–1190. doi:10.1016/j.bbadis.2016.03.007

6. Bonacina F, Moregola A, Porte R, Baragetti A, Bonavita E, Salatin A, Grigore L, Pellegatta F, Molgora M, Sironi M, Barbati E, Mantovani A, Bottazzi B, Catapano AL, Garlanda C, Norata GD. 2019. Pentraxin 3 deficiency protects from the metabolic inflammation associated to diet-induced obesity. Cardiovascular Research 115:1861–1872. doi:10.1093/cvr/cvz068

7. Bottazzi B, Doni A, Garlanda C, Mantovani A. 2010. An Integrated View of Humoral Innate Immunity: Pentraxins as a Paradigm. Annu Rev Immunol 28:157–183. doi:10.1146/annurev-immunol-030409-101305

8. Bottazzi B, Santini L, Savino S, Giuliani MM, Dueñas Díez AI, Mancuso G, Beninati C, Sironi M, Valentino S, Deban L, Garlanda C, Teti G, Pizza M, Rappuoli R, Mantovani A. 2015. Recognition of Neisseria meningitidis by the Long Pentraxin PTX3 and Its Role as an Endogenous Adjuvant. PLoS ONE 10:e0120807. doi:10.1371/journal.pone.0120807

9. Bottazzi B, Vouret-Craviari V, Bastone A, De Gioia L, Matteucci C, Peri G, Spreafico F, Pausa M, D’Ettorre C, Gianazza E, Tagliabue A, Salmona M, Tedesco F, Introna M, Mantovani A. 1997. Multimer Formation and Ligand Recognition by the Long Pentraxin PTX3. Journal of Biological Chemistry 272:32817–32823. doi:10.1074/jbc.272.52.32817

10. Bou Ghanem EN, Clark S, Roggensack SE, McIver SR, Alcaide P, Haydon PG, Leong JM. 2015. Extracellular Adenosine Protects against Streptococcus pneumoniae Lung Infection by Regulating Pulmonary Neutrophil Recruitment. PLoS Pathog 11:e1005126. doi:10.1371/journal.ppat.1005126

11. Brunel A-S, Wójtowicz A, Lamoth F, Spertini O, Neofytos D, Calandra T, Marchetti O, Bochud P-Y. 2018. Pentraxin-3 polymorphisms and invasive mold infections in acute leukemia patients receiving intensive chemotherapy. Haematologica 103:e527–e530. doi:10.3324/haematol.2018.195453

12. Campos CF, Leite L, Pereira P, Vaz CP, Branca R, Campilho F, Freitas F, Ligeiro D, Marques A, Torrado E, Silvestre R, Lacerda JF, Campos Jr. A, Cunha C, Carvalho A. 2019. PTX3 Polymorphisms Influence Cytomegalovirus Reactivation After Stem-Cell Transplantation. Front Immunol 10:88. doi:10.3389/fimmu.2019.00088

13. Chiarini M, Sabelli C, Melotti P, Garlanda C, Savoldi G, Mazza C, Padoan R, Plebani A, Mantovani A, Notarangelo LD, Assael BM, Badolato R. 2010. PTX3 genetic variations affect the risk of Pseudomonas aeruginosa airway colonization in cystic fibrosis patients. Genes Immun 11:665–670. doi:10.1038/gene.2010.41

14. Cunha C, Aversa F, Lacerda JF, Busca A, Kurzai O, Grube M, Löffler J, Maertens JA, Bell AS, Inforzato A, Barbati E, Almeida B, Santose Sousa P, Barbui A, Potenza L, Caira M, Rodrigues F, Salvatori G, Pagano L, Luppi M, Mantovani A, Velardi A, Romani L, Carvalho A. 2014. Genetic PTX3 Deficiency and Aspergillosis in Stem-Cell Transplantation. N Engl J Med 370:421–432. doi:10.1056/NEJMoa1211161

15. Cunha C, Monteiro AA, Oliveira-Coelho A, Kühne J, Rodrigues F, Sasaki SD, Schio SM, Camargo JJ, Mantovani A, Carvalho A, Pasqualotto AC. 2015. PTX3-Based Genetic Testing for Risk of Aspergillosis After Lung Transplant: Table 1. Clin Infect Dis 61:1893–1894. doi:10.1093/cid/civ679

16. Daigo K, Yamaguchi N, Kawamura T, Matsubara K, Jiang S, Ohashi R, Sudou Y, Kodama T, Naito M, Inoue K, Hamakubo T. 2012. The Proteomic Profile of Circulating Pentraxin 3 (PTX3) Complex in Sepsis Demonstrates the Interaction with Azurocidin 1 and Other Components of Neutrophil Extracellular Traps. Molecular & Cellular Proteomics 11:M111.015073. doi:10.1074/mcp.M111.015073

17. de Porto AP, Liu Z, de Beer R, Florquin S, de Boer OJ, Hendriks RW, van der Poll T, de Vos AF. 2019. Btk inhibitor ibrutinib reduces inflammatory myeloid cell responses in the lung during murine pneumococcal pneumonia. Mol Med 25:3. doi:10.1186/s10020-018-0069-7

18. Deban L, Russo RC, Sironi M, Moalli F, Scanziani M, Zambelli V, Cuccovillo I, Bastone A, Gobbi M, Valentino S, Doni A, Garlanda C, Danese S, Salvatori G, Sassano M, Evangelista V, Rossi B, Zenaro E, Constantin G, Laudanna C, Bottazzi B, Mantovani A. 2010. Regulation of leukocyte recruitment by the long pentraxin PTX3. Nat Immunol 11:328–334. doi:10.1038/ni.1854

19. Doni A, Musso T, Morone D, Bastone A, Zambelli V, Sironi M, Castagnoli C, Cambieri I, Stravalaci M, Pasqualini F, Laface I, Valentino S, Tartari S, Ponzetta A, Maina V, Barbieri SS, Tremoli E, Catapano AL, Norata GD, Bottazzi B, Garlanda C, Mantovani A. 2015. An acidic microenvironment sets the humoral pattern recognition molecule PTX3 in a tissue repair mode 212:21.

20. Faul F, Erdfelder E, Lang A-G, Buchner A. 2007. G*Power 3: a flexible statistical power analysis program for the social, behavioral, and biomedical sciences. Behav Res Methods 39:175–191. doi:10.3758/bf03193146

21. García-Laorden MI, Hernández-Brito E, Muñoz-Almagro C, Pavlovic-Nesic S, Rúa-Figueroa I, Briones ML, Rajas O, Borderías L, Payeras A, Lorente L, Freixinet J, Ferreres J, Obando I, González-Quevedo N, Rodríguez de Castro F, Solé-Violán J, Rodríguez-Gallego C. 2020. Should MASP-2 Deficiency Be Considered a Primary Immunodeficiency? Relevance of the Lectin Pathway. J Clin Immunol 40:203–210. doi:10.1007/s10875-019-00714-4

22. Garlanda C, Bottazzi B, Magrini E, Inforzato A, Mantovani A. 2018. PTX3, a Humoral Pattern Recognition Molecule, in Innate Immunity, Tissue Repair, and Cancer. Physiological Reviews 98:623–639. doi:10.1152/physrev.00016.2017

23. Garlanda C, Hirsch E, Bozza S, Salustri A, De Acetis M, Nota R, Maccagno A, Riva F, Bottazzi B, Peri G, Doni A, Vago L, Botto M, De Santis R, Carminati P, Siracusa G, Altruda F, Vecchi A, Romani L, Mantovani A. 2002. Non-redundant role of the long pentraxin PTX3 in anti-fungal innate immune response. Nature 420:182–186. doi:10.1038/nature01195

24. Haapasalo K, Meri S. 2019. Regulation of the Complement System by Pentraxins. Front Immunol 10:1750. doi:10.3389/fimmu.2019.01750

25. Hasenberg M, Behnsen J, Krappmann S, Brakhage A, Gunzer M. 2011. Phagocyte responses towards Aspergillus fumigatus. International Journal of Medical Microbiology 301:436–444. doi:10.1016/j.ijmm.2011.04.012

26. He Q, Li H, Rui Y, Liu L, He B, Shi Y, Su X. 2018. Pentraxin 3 Gene Polymorphisms and Pulmonary Aspergillosis in Chronic Obstructive Pulmonary Disease Patients. Clinical Infectious Diseases 66:261–267. doi:10.1093/cid/cix749

27. Hommes TJ, Hoogendijk AJ, Dessing MC, Van’t Veer C, Florquin S, Colonna M, de Vos AF, van der Poll T. 2014. Triggering receptor expressed on myeloid cells-1 (TREM-1) improves host defence in pneumococcal pneumonia. J Pathol 233:357–367. doi:10.1002/path.4361

28. Inforzato A, Rivieccio V, Morreale AP, Bastone A, Salustri A, Scarchilli L, Verdoliva A, Vincenti S, Gallo G, Chiapparino C, Pacello L, Nucera E, Serlupi-Crescenzi O, Day AJ, Bottazzi B, Mantovani A, De Santis R, Salvatori G. 2008. Structural characterization of PTX3 disulfide bond network and its multimeric status in cumulus matrix organization. J Biol Chem 283:10147–10161. doi:10.1074/jbc.M708535200

29. Jaillon S, Moalli F, Ragnarsdottir B, Bonavita E, Puthia M, Riva F, Barbati E, Nebuloni M, Cvetko Krajinovic L, Markotic A, Valentino S, Doni A, Tartari S, Graziani G, Montanelli A, Delneste Y, Svanborg C, Garlanda C, Mantovani A. 2014. The Humoral Pattern Recognition Molecule PTX3 Is a Key Component of Innate Immunity against Urinary Tract Infection. Immunity 40:621–632. doi:10.1016/j.immuni.2014.02.015

30. Jaillon S, Peri G, Delneste Y, Frémaux I, Doni A, Moalli F, Garlanda C, Romani L, Gascan H, Bellocchio S, Bozza S, Cassatella MA, Jeannin P, Mantovani A. 2007. The humoral pattern recognition receptor PTX3 is stored in neutrophil granules and localizes in extracellular traps. Journal of Experimental Medicine 204:793–804. doi:10.1084/jem.20061301

31. Kao S-J, Yang H-W, Tsao S-M, Cheng C-W, Bien M-Y, Yu M-C, Bai K-J, Yang S-F, Chien M-H. 2013. Plasma long pentraxin 3 (PTX3) concentration is a novel marker of disease activity in patients with community-acquired pneumonia. Clinical Chemistry and Laboratory Medicine 51. doi:10.1515/cclm-2012-0459

32. Kjos M, Aprianto R, Fernandes VE, Andrew PW, van Strijp JAG, Nijland R, Veening J-W. 2015. Bright Fluorescent Streptococcus pneumoniae for Live-Cell Imaging of Host-Pathogen Interactions. J Bacteriol 197:807–818. doi:10.1128/JB.02221-14

33. Koh SH, Shin SG, Andrade MJ, Go R-H, Park S, Woo C-H, Lim JH. 2017. Long pentraxin PTX3 mediates acute inflammatory responses against pneumococcal infection. Biochem Biophys Res Commun 493:671–676. doi:10.1016/j.bbrc.2017.08.133

34. Lakens D. 2013. Calculating and reporting effect sizes to facilitate cumulative science: a practical primer for t-tests and ANOVAs. Front Psychol 4:863. doi:10.3389/fpsyg.2013.00863

35. Lech M, Römmele C, Gröbmayr R, Eka Susanti H, Kulkarni OP, Wang S, Gröne H-J, Uhl B, Reichel C, Krombach F, Garlanda C, Mantovani A, Anders H-J. 2013. Endogenous and exogenous pentraxin-3 limits postischemic acute and chronic kidney injury. Kidney International 83:647–661. doi:10.1038/ki.2012.463

36. Liu F, Wu HY, Wesselschmidt R, Kornaga T, Link DC. 1996. Impaired Production and Increased Apoptosis of Neutrophils in Granulocyte Colony-Stimulating Factor Receptor–Deficient Mice. Immunity 5:491–501. doi:10.1016/S1074-7613(00)80504-X

37. Lovewell RR, Patankar YR, Berwin B. 2014. Mechanisms of phagocytosis and host clearance of *Pseudomonas aeruginosa*. American Journal of Physiology-Lung Cellular and Molecular Physiology 306:L591–L603. doi:10.1152/ajplung.00335.2013

38. Maas SL, Soehnlein O, Viola JR. 2018. Organ-Specific Mechanisms of Transendothelial Neutrophil Migration in the Lung, Liver, Kidney, and Aorta. Front Immunol 9:2739. doi:10.3389/fimmu.2018.02739

39. Madouri F, Barada O, Kervoaze G, Trottein F, Pichavant M, Gosset P. 2018. Production of Interleukin-20 cytokines limits bacterial clearance and lung inflammation during infection by Streptococcus pneumoniae. EBioMedicine 37:417–427. doi:10.1016/j.ebiom.2018.10.031

40. Mauri T, Coppadoro A, Bombino M, Bellani G, Zambelli V, Fornari C, Berra L, Bittner EA, Schmidt U, Sironi M, Bottazzi B, Brambilla P, Mantovani A, Pesenti A. 2014. Alveolar pentraxin 3 as an early marker of microbiologically confirmed pneumonia: a threshold-finding prospective observational study. Crit Care 18:562. doi:10.1186/s13054-014-0562-5

41. Moalli F, Doni A, Deban L, Zelante T, Zagarella S, Bottazzi B, Romani L, Mantovani A, Garlanda C. 2010. Role of complement and Fc{gamma} receptors in the protective activity of the long pentraxin PTX3 against Aspergillus fumigatus. Blood 116:11. doi:10.1182/blood-2009-12-258376

42. Moalli F, Paroni M, Véliz Rodriguez T, Riva F, Polentarutti N, Bottazzi B, Valentino S, Mantero S, Nebuloni M, Mantovani A, Bragonzi A, Garlanda C. 2011. The Therapeutic Potential of the Humoral Pattern Recognition Molecule PTX3 in Chronic Lung Infection Caused by *Pseudomonas aeruginosa*. JI 186:5425– 5434. doi:10.4049/jimmunol.1002035

43. Nathan C. 2006. Neutrophils and immunity: challenges and opportunities. Nat Rev Immunol 6:173–182. doi:10.1038/nri1785

44. Olesen R, Wejse C, Velez DR, Bisseye C, Sodemann M, Aaby P, Rabna P, Worwui A, Chapman H, Diatta M, Adegbola RA, Hill PC, Østergaard L, Williams SM, Sirugo G. 2007. DC-SIGN (CD209), pentraxin 3 and vitamin D receptor gene variants associate with pulmonary tuberculosis risk in West Africans. Genes Immun 8:456–467. doi:10.1038/sj.gene.6364410

45. Ponzetta A, Carriero R, Carnevale S, Barbagallo M, Molgora M, Perucchini C, Magrini E, Gianni F, Kunderfranco P, Polentarutti N, Pasqualini F, Di Marco S, Supino D, Peano C, Cananzi F, Colombo P, Pilotti S, Alomar SY, Bonavita E, Galdiero MR, Garlanda C, Mantovani A, Jaillon S. 2019. Neutrophils Driving Unconventional T Cells Mediate Resistance against Murine Sarcomas and Selected Human Tumors. Cell 178:346–360.e24. doi:10.1016/j.cell.2019.05.047

46. Porte R, Davoudian S, Asgari F, Parente R, Mantovani A, Garlanda C, Bottazzi B. 2019. The Long Pentraxin PTX3 as a Humoral Innate Immunity Functional Player and Biomarker of Infections and Sepsis. Front Immunol 10:794. doi:10.3389/fimmu.2019.00794

47. Porte R, Fougeron D, Muñoz-Wolf N, Tabareau J, Georgel A-F, Wallet F, Paget C, Trottein F, Chabalgoity JA, Carnoy C, Sirard J-C. 2015. A Toll-Like Receptor 5 Agonist Improves the Efficacy of Antibiotics in Treatment of Primary and Influenza Virus-Associated Pneumococcal Mouse Infections. Antimicrob Agents Chemother 59:6064–6072. doi:10.1128/AAC.01210-15

48. Purcell S, Neale B, Todd-Brown K, Thomas L, Ferreira MAR, Bender D, Maller J, Sklar P, de Bakker PIW, Daly MJ, Sham PC. 2007. PLINK: A Tool Set for Whole-Genome Association and Population-Based Linkage Analyses. The American Journal of Human Genetics 81:559–575. doi:10.1086/519795

49. Quinn MT, DeLeo FR, editors. 2020. Neutrophil: Methods and Protocols, Methods in Molecular Biology. New York, NY: Springer US. doi:10.1007/978-1-0716-0154-9

50. Quinton LJ, Mizgerd JP. 2015. Dynamics of Lung Defense in Pneumonia: Resistance, Resilience, and Remodeling. Annu Rev Physiol 77:407–430. doi:10.1146/annurev-physiol-021014-071937

51. Saleh MAA, van de Garde EMW, van Hasselt JGC. 2019. Host-response biomarkers for the diagnosis of bacterial respiratory tract infections. Clinical Chemistry and Laboratory Medicine (CCLM) 57:442–451. doi:10.1515/cclm-2018-0682

52. Schouten M, de Boer JD, Kager LM, Roelofs JJTH, Meijers JCM, Esmon CT, Levi M, van’t Veer C, van der Poll T. 2014. The endothelial protein C receptor impairs the antibacterial response in murine pneumococcal pneumonia and sepsis. Thromb Haemost 111:970–980. doi:10.1160/TH13-10-0859

53. Schwab S, Jobin K, Kurts C. 2017. Urinary tract infection: recent insight into the evolutionary arms race between uropathogenic Escherichia coli and our immune system. Nephrology Dialysis Transplantation 32:1977–1983. doi:10.1093/ndt/gfx022

54. Sender V, Hentrich K, Pathak A, Tan Qian Ler A, Embaie BT, Lundström SL, Gaetani M, Bergstrand J, Nakamoto R, Sham L-T, Widengren J, Normark S, Henriques-Normark B. 2020. Capillary leakage provides nutrients and antioxidants for rapid pneumococcal proliferation in influenza-infected lower airways. Proc Natl Acad Sci U S A 117:31386–31397. doi:10.1073/pnas.2012265117

55. Shi G-Q, Yang L, Shan L-Y, Yin L-Z, Jiang W, Tian H-T, Yang D-D. 2020. Investigation of the clinical significance of detecting PTX3 for community-acquired pneumonia. European Review for Medical and Pharmacological Sciences 24:8477–8482. doi:10.26355/eurrev_202008_22645

56. Siljan WW, Holter JC, Michelsen AE, Nymo SH, Lauritzen T, Oppen K, Husebye E, Ueland T, Mollnes TE, Aukrust P, Heggelund L. 2019. Inflammatory biomarkers are associated with aetiology and predict outcomes in community-acquired pneumonia: results of a 5-year follow-up cohort study. ERJ Open Res 5:00014–02019. doi:10.1183/23120541.00014-2019

57. Sohail I, Ghosh S, Mukundan S, Zelewski S, Khan MN. 2018. Role of Inflammatory Risk Factors in the Pathogenesis of Streptococcus pneumoniae. Front Immunol 9:2275. doi:10.3389/fimmu.2018.02275

58. Stravalaci M, Pagani I, Paraboschi EM, Pedotti M, Doni A, Scavello F, Mapelli SN, Sironi M, Perucchini C, Varani L, Matkovic M, Cavalli A, Cesana D, Gallina P, Pedemonte N, Capurro V, Clementi N, Mancini N, Invernizzi P, Bayarri-Olmos R, Garred P, Rappuoli R, Duga S, Bottazzi B, Uguccioni M, Asselta R, Vicenzi E, Mantovani A, Garlanda C. 2022. Recognition and inhibition of SARS-CoV-2 by humoral innate immunity pattern recognition molecules. Nat Immunol 23:275–286. doi:10.1038/s41590-021-01114-w

59. Szalai AJ. 2002. The antimicrobial activity of C-reactive protein. Microbes and Infection 4:201–205. doi:10.1016/S1286-4579(01)01528-3

60. Tavares LP, Garcia CC, Vago JP, Queiroz-Junior CM, Galvão I, David BA, Rachid MA, Silva PMR, Russo RC, Teixeira MM, Sousa LP. 2016. Inhibition of Phosphodiesterase-4 during Pneumococcal Pneumonia Reduces Inflammation and Lung Injury in Mice. Am J Respir Cell Mol Biol 55:24–34. doi:10.1165/rcmb.2015-0083OC

61. Thulborn SJ, Dilpazir M, Haldar K, Mistry V, Brightling CE, Barer MR, Bafadhel M. 2017. Investigating the role of pentraxin 3 as a biomarker for bacterial infection in subjects with COPD. COPD **Volume** 12:1199–1205. doi:10.2147/COPD.S123528

62. Tin Tin Htar M, Morato Martínez J, Theilacker C, Schmitt H-J, Swerdlow D. 2019. Serotype evolution in Western Europe: perspectives on invasive pneumococcal diseases (IPD). Expert Review of Vaccines 18:1145–1155. doi:10.1080/14760584.2019.1688149

63. Troeger C, Blacker B, Khalil IA, Rao PC, Cao J, Zimsen SRM, Albertson SB, Deshpande A, Farag T, Abebe Z, Adetifa IMO, Adhikari TB, Akibu M, Al Lami FH, Al-Eyadhy A, Alvis-Guzman N, Amare AT, Amoako YA, Antonio CAT, Aremu O, Asfaw ET, Asgedom SW, Atey TM, Attia EF, Avokpaho EFGA, Ayele HT, Ayuk TB, Balakrishnan K, Barac A, Bassat Q, Behzadifar Masoud, Behzadifar Meysam, Bhaumik S, Bhutta ZA, Bijani A, Brauer M, Brown A, Camargos PAM, Castañeda-Orjuela CA, Colombara D, Conti S, Dadi AF, Dandona L, Dandona R, Do HP, Dubljanin E, Edessa D, Elkout H, Endries AY, Fijabi DO, Foreman KJ, Forouzanfar MH, Fullman N, Garcia-Basteiro AL, Gessner BD, Gething PW, Gupta R, Gupta T, Hailu GB, Hassen HY, Hedayati MT, Heidari M, Hibstu DT, Horita N, Ilesanmi OS, Jakovljevic MB, Jamal AA, Kahsay A, Kasaeian A, Kassa DH, Khader YS, Khan EA, Khan MN, Khang Y-H, Kim YJ, Kissoon N, Knibbs LD, Kochhar S, Koul PA, Kumar GA, Lodha R, Magdy Abd El Razek H, Malta DC, Mathew JL, Mengistu DT, Mezgebe HB, Mohammad KA, Mohammed MA, Momeniha F, Murthy S, Nguyen CT, Nielsen KR, Ningrum DNA, Nirayo YL, Oren E, Ortiz JR, Pa M, Postma MJ, Qorbani M, Quansah R, Rai RK, Rana SM, Ranabhat CL, Ray SE, Rezai MS, Ruhago GM, Safiri S, Salomon JA, Sartorius B, Savic M, Sawhney M, She J, Sheikh A, Shiferaw MS, Shigematsu M, Singh JA, Somayaji R, Stanaway JD, Sufiyan MB, Taffere GR, Temsah M-H, Thompson MJ, Tobe-Gai R, Topor-Madry R, Tran BX, Tran TT, Tuem KB, Ukwaja KN, Vollset SE, Walson JL, Weldegebreal F, Werdecker A, West TE, Yonemoto N, Zaki MES, Zhou L, Zodpey S, Vos T, Naghavi M, Lim SS, Mokdad AH, Murray CJL, Hay SI, Reiner RC. 2018. Estimates of the global, regional, and national morbidity, mortality, and aetiologies of lower respiratory infections in 195 countries, 1990–2016: a systematic analysis for the Global Burden of Disease Study 2016. The Lancet Infectious Diseases 18:1191–1210. doi:10.1016/S1473-3099(18)30310-4

64. Weinberger DM, Harboe ZB, Sanders EAM, Ndiritu M, Klugman KP, Rückinger S, Dagan R, Adegbola R, Cutts F, Johnson HL, O’Brien KL, Anthony Scott J, Lipsitch M. 2010. Association of Serotype with Risk of Death Due to Pneumococcal Pneumonia: A Meta-Analysis. Clin Infect Dis 51:692–699. doi:10.1086/655828

65. Weiser JN, Ferreira DM, Paton JC. 2018. Streptococcus pneumoniae: transmission, colonization and invasion. Nat Rev Microbiol 16:355–367. doi:10.1038/s41579-018-0001-8

66. Wigginton JE, Cutler DJ, Abecasis GR. 2005. A Note on Exact Tests of Hardy-Weinberg Equilibrium. The American Journal of Human Genetics 76:887–893. doi:10.1086/429864

67. Wójtowicz A, Lecompte TD, Bibert S, Manuel O, Rüeger S, Berger C, Boggian K, Cusini A, Garzoni C, Hirsch H, Khanna N, Mueller NJ, Meylan PR, Pascual M, van Delden C, Bochud P-Y. 2015. *PTX3* Polymorphisms and Invasive Mold Infections After Solid Organ Transplant: Figure 1. Clin Infect Dis 61:619–622. doi:10.1093/cid/civ386

